# BEN-solo factors partition active chromatin to ensure proper gene activation in *Drosophila*

**DOI:** 10.1101/593830

**Authors:** Malin Ueberschär, Huazhen Wang, Chun Zhang, Shu Kondo, Tsutomu Aoki, Paul Schedl, Eric C. Lai, Jiayu Wen, Qi Dai

## Abstract

The *Drosophila* genome encodes three BEN-solo proteins including Insensitive (Insv), Elba1 and Elba2 that possess activities in both transcriptional repression and chromatin insulation. These proteins all have a DNA binding BEN domain. A fourth protein Elba3 bridges Elba1 and Elba2 to form a heterotrimeric complex ELBA. Here we report comprehensive investigation on the *in vivo* functions of these proteins in *Drosophila* embryos. We generate ChIP-seq data of all these factors from all cognate and non-cognate mutants to assess common and distinct binding locations of Insv and ELBA, and genetic interdependencies. Notably, while most Elba1 and Elba2 binding requires the full ELBA complex, the adapter protein Elba3 can associate with chromatin and repress gene expression independently of Elba1 and Elba2. We also employ high-resolution ChIP-nexus mapping to show that Insv binds to DNA in a symmetric configuration while the ELBA complex binds asymmetrically *in vivo*. We observe that motifs of known insulator proteins are enriched in ELBA and Insv ChIP peaks and demonstrate that ELBA collaborates with other insulator factors to regulate developmental patterning in embryos. To differentiate the insulator function of ELBA and Insv from their repressor activity, we determined real-time transcription change in mutant embryos using precision nuclear run-on sequencing. ELBA factor mutants dampen expression differences between pairs of ELBA-bound neighboring genes. Finally, transgenic reporters confirm insulation activity of ELBA- and Insv-bound sites. Altogether, these findings define ELBA and Insv as general insulator proteins in *Drosophila* and demonstrate the functional importance of insulators in partitioning transcription units.

## Introduction

Proper gene regulation requires coordinated activities of distinct classes of cis- and trans-regulators. Insulators (or boundary elements) are a special type of cis elements that constrain enhancer-promoter interactions (Holdridge and Dorsett 1991; Geyer and Corces 1992; Kellum and Schedl 1992; Dorsett 1993; Bell et al. 1999) and set chromatin boundaries (Kellum and Schedl 1991). Historically, boundary or enhancer-blocking activities of newly identified insulators were mostly tested on a one-on-one basis in transgenic lines or genetically dissected for individual loci. Recent advances in genomics and chromatin structure capture techniques allowed more systematic identification of insulators and but also assigned new properties to them in chromatin architecture organization (reviewed in (Phillips-Cremins and Corces 2013), (Valenzuela and Kamakaka 2006)).

The activity of insulator elements is realized via associated insulator factors. The zinc-finger protein CTCF seems to be the only insulator protein conserved between vertebrates and invertebrates. In addition to its established roles as an insulator in chromatin organization, long-range regulatory element looping and enhancer segregating (Phillips-Cremins and Corces 2013), several original studies on mammalian CTCF indicated its direct role in transcriptional repression (Lutz et al. 2000; Perez-Juste et al. 2000). In contrast, more than a dozen proteins were indicated with insulator function in *Drosophila* (Kyrchanova and Georgiev 2014). According to the binding patterns bound by five classic insulators including CP190, BEAF32, CTCF, Su(Hw) and Mod(mdg4), *Drosophila* insulators were divided into two classes (Negre et al. 2010). Class I are mainly bound by CP190, BEAF32 and CTCF in active chromatin regions proximal to promoters, while class II insulators are mostly bound by Su(Hw) located in more distal intergenic loci. However, at functional level, how these factors cooperate with each other remains unclear.

The BEN (BANP, E5R, and NAC1) domain is a recently recognized domain present in a variety of metazoan and viral proteins (Abhiman et al. 2008). Several BEN-containing proteins including mammalian BANP/SMAR1 (Kaul-Ghanekar et al. 2004; Rampalli et al. 2005), NAC1 (Korutla et al. 2005; Korutla et al. 2007), BEND3 (Sathyan et al. 2011), and the C isoform of *Drosophila* mod(mdg4) (Gerasimova et al. 1995; Negre et al. 2010) have chromatin associated function and have been linked to transcriptional silencing. We and others showed that the BEN domain possesses an intrinsic sequence-specific DNA binding activity. Mammalian RBB, a BEN and BTB domain protein binds to and directly represses expression of the HDM2 oncogene through interacting with the nucleosome remodeling and deacetylase (NuRD) complex (Xuan et al. 2013). *Drosophila* Insv binds to a palindromic motif, TCCAATTGGA and its variants (TCYAATHRGAA), and represses genes in the nervous system (Dai et al. 2013b). Two other *Drosophila* BEN proteins, Elba1 and Elba2, along with the adaptor protein Elba3, are assembled in a hetero-trimeric complex (ELBA) and associate with the asymmetric site “CCAATAAG” in the Fab-7 insulator (Aoki et al. 2012). The closely linked *elba1* and *elba3* genes are specifically expressed during the mid-blastula transition, which restricts ELBA activity to this early developmental window. Interestingly, the genes encoding Insv and Elba2 are also arranged head-to-head in the genome, while their gene products are present throughout development.

Most of BEN domain proteins contain other characterized motifs. However, Insv, Elba1, Elba2 and several mammalian homologs, such as BEND5 and BEND6, harbor only one BEN domain and lack other known functional domains. Thus, we referr to this sub-class as BEN-solo factors (Dai et al. 2013a; Dai et al. 2015). Our previous work suggests that Insv and ELBA BEN-solo factors share common properties, *e.g*. binding to the palindromic sites as homodimers and repressing reporter genes in culture cells, but also display distinct activities, *e.g*. Insv being the only one that interacts with Notch signaling in *Drosophila* peripheral nervous system and its inability to bind to the asymmetric site (Dai et al. 2015). Interestingly, the Fab-7 insulator requires ELBA for its early boundary activity but also requires Insv in later development (Fedotova et al. 2018).

It remains to be determined how the ELBA factors regulate gene expression and what the biological functions of ELBA and Insv in embryogenesis are. In this study, we comprehensively characterized the three *Drosophila* BEN-solo factors and the adapter protein Elba3 in early *Drosophila* embryos, by analyzing mutants, their DNA binding preferences (symmetric versus asymmetric), chromatin binding inter-dependence (homo-dimers versus hetero-trimeric complex) and mechanisms in gene regulation (repressor versus insulator). Our ChIP-seq analyses confirm that all three BEN-solo factors associate with both the symmetric (palindromic) and asymmetric types of sites, despite that Insv displays higher affinity to the symmetric type. Unexpectedly, Elba3 remains associated with chromatin even in the absence of its DNA binding partners Elba1 and Elba2, suggesting that it also uses other co-factor(s) to target chromatin. Our ChIP-nexus (chromatin immunoprecipitation experiments with nucleotide resolution through exonuclease, unique barcode and single ligation) assay shows a genome-wide symmetric DNA binding pattern of Insv in contrast to an asymmetric pattern of Elba1 and Elba2. Consistent with the repressive function in culture cells, their direct targets became de-repressed in mutant embryos. Moreover, the ELBA factors and the other insulator proteins including GAF and CP190 show strong genetic interactions in regulating embryonic patterning. Finally, using the PRO-seq approach, we show that adjacent genes flanked with ELBA binding are less differentially expressed in all three *ELBA* mutants. Insv-associated adjacent genes do not show such a global effect, but individual neighbor promoters are insulated by Insv binding. These findings indicate a role of ELBA and Insv as general insulators in partitioning transcription units in *Drosophila*. In support of this conclusion, ELBA- and Insv-bound elements show blocking of enhancer-promoter interaction in transgenic reporters.

## RESULTS

### The ELBA complex shares many genomic binding sites with Insv

We previously described genomic binding for Insv whose ChIP-seq peaks cover numerous genomic sites that bear its specific binding motif (CCAATTGG and variants thereof) (Dai et al. 2013a). A few individual sites in the Fab-7 and the Fab-8 insulators were known genomic locations of the ELBA complex (Aoki et al. 2012; Aoki et al. 2014; Fedotova et al. 2018). We intended to broaden this perspective by generating ChIP-seq data for each of the three ELBA factors from the blastoderm stage of embryos, which covers the peak expression of the ELBA factors (Dai et al. 2015) (**Supplemental Fig.S1A**). We also made frame-shift mutant alleles for *elba1*, *elba2* and *elba3* by using a guide RNA targeting each gene locus with CRISPR/Cas9. The mutant animals of the ELBA factors are viable, similar to *insv* mutant, making it possible to obtain sufficient ChIP-seq material from mutants of the same stage (**Supplemental table 1**). For uniformity of comparing peak-calls, we re-made Insv ChIP-seq data from the same stage of embryos in parallel with the ELBA libraries.

To determine ELBA and Insv binding regions, we first assessed the quality of the three controls, Input, IgG and mutant ChIP. Mutant ChIP data appeared to be the most stringent, as it gave the highest occurrence of the known Insv/ELBA motif (**Supplemental Fig. S1B**). We called 3151, 1468, 6525 and 4927 peaks for Elba1, Elba2, Elba3 and Insv respectively after using wild-type ChIP-seq peaks against the cognate mutant ChIP peaks (Fig. 1 A-B**, Supplemental table2**). When ranked according to the peak scores of the Elba3 ChIP signal, the three ELBA peaks show extensive correlation, whereas a group of strong Insv peaks stand out and do not correlate with the ELBA peaks (Fig. 1 A). Similarly, the Venn diagram shows that about half of the Insv peaks are unique (Fig. 1 B), while the Elba2 peaks are covered entirely by the Elba1 peaks that are further covered by the Elba3 peaks. We reasoned that the difference in the number of the three ELBA ChIP peaks could be due to difference on their antibody affinities or in the number of *in vivo* binding regions. Indeed, the Elba2 antibody did not work well in immunofluorescent staining whereas the Elba3 antibody showed strongest signal (**Supplemental Fig. S1A**).

**Figure 1:**
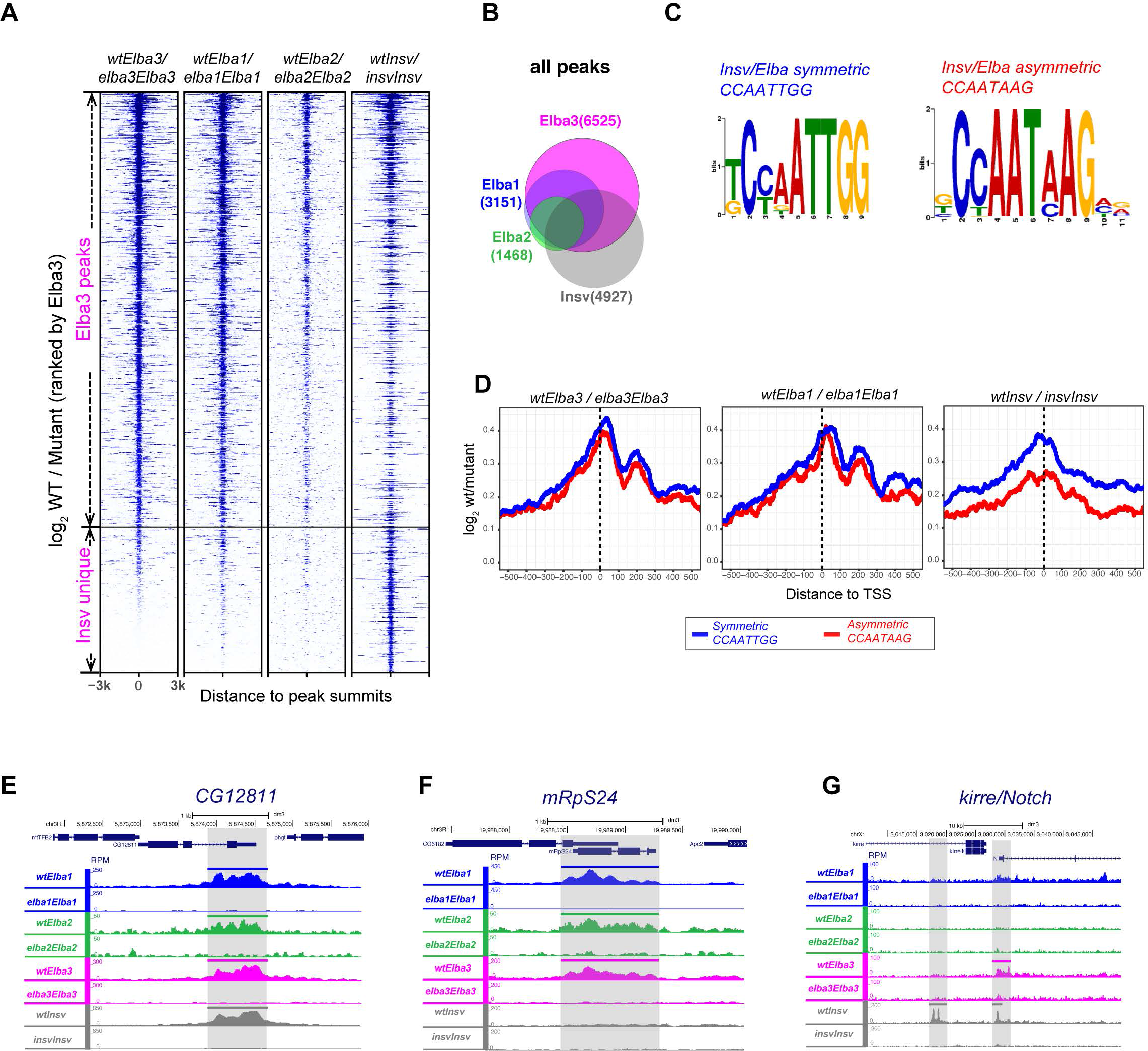
Comparison of the ELBA complex with Insv binding sites and motifs. (**A**) A heatmap of ChIP-seq coverage ratio (log2 wt/mutant) centered at peak summits for the four Elba/Insv factors ranked by the Elba3 signal, showing Elba1, Elba2, and Elba3 peaks largely overlap while Insv has unique peaks. The gene names with all the letters in lower case denote genotypes and the names with the first letter upper-case denote antibodies in ChIP. For each of the four factors, the peaks were called by using the ChIP-seq reads of wt against the ChIP reads of its cognate mutant. (**B**) A Venn diagram shows overlapping fractions for four factors. (**C**) Logos of the Insv symmetric and the ELBA-type asymmetric motifs from *de novo* motif discovery. (**D**) The coverage ratio (log2 wt/mutant) centered at TSS is shown for the peaks that contain either the symmetric or asymmetric motif. The symmetric and asymmetric motifs are equally enriched in the Elba peaks while the symmetric motif has a higher enrichment than the asymmetric motif in the Insv peaks. (**E-F**) Screenshots of three example loci: all four factors bound sites in *CG12811* (**E**), Elba1/2/3 unique sites in *mRpS24* (**F**), and Insv unique site in *Kirre/Notch* (**G**). The coverage tracks were normalized to the library sizes to give Read Per Million (RPM) per base.

Our *de novo* motif discovery analysis identified the known Insv symmetric CCAATTGG and the ELBA-type asymmetric sites CCAATAAG as well as their variants from the ChIP-seq peaks of all four factors (Fig. 1 C-D). The two types of motifs are similarly enriched in the ELBA factor peaks, consistent with our previous report that Elba1 and Elba2 can bind to both motifs *in vitro*. For Insv, the peaks that contain the symmetric sites show higher ChIP signal than those with the asymmetric ones (Fig. 1 D), suggesting that Insv binds to the symmetric site with higher affinity or in more loci.

Compared to the Elba3 peaks that enrich at promoter proximal regions, a larger fraction of the Insv-unique sites locates at distal upstream regions, introns, exons and intergenic regions (**Supplemental Fig. S1C**). The Insv-unique portion also covers more peaks that contain the consensus sequences (**Supplemental Fig. S1D**). Thus, despite having similar DNA binding domains and expression patterns in early embryos, Insv displays distinct binding preferences compared to ELBA. Three loci exemplifying four-factor binding (*CG12811*), ELBA unique binding (*mRpS24*) and Insv unique binding (*kirre/Notch*) are illustrated (Fig. 1E-G). Notably, the Insv-unique peak locates in the intronic region of *kirre* and upstream of the *Notch* locus (Fig. 1G). The Elba3/Insv peak at the upstream of the *Notch* locus covers a known insulator element, facet-strawberry (Vazquez and Schedl 2000). As Insv is the only factor that shows interaction with the Notch pathway in sensory organ development (Duan et al. 2011; Dai et al. 2015), it will be of interest to know whether the Insv sites in this region regulate expression of *Notch*.

### Elba1 and Elba3 are able to associate with chromatin independent of the ELBA trimeric complex

The three subunits of ELBA rely on one another to be able to bind to the ELBA site in Fab-7 *in vitro* (Aoki et al. 2012). We generated ChIP-seq data for each factor from each non-cognate mutant background (Fig. 2A, **Supplemental table 2**). This allowed us to assess possible genetic inter-dependencies of binding amongst Insv and ELBA factors. Interestingly, binding of Elba1 was completely lost in *elba3* mutant, and binding of Elba2 was nearly eliminated in this background (with only 68 peaks remained). This was unexpected since our previous ectopic expression experiments indicated that Elba1 and Elba2 can form homodimers in cultured cells (Dai et al. 2015). By contrast, this data indicates that, in the early embryo, Elba3 is an essential component for the endogenous ELBA complex to bind the genome. Most of the Elba1 binding sites are lost in *elba2* mutant with only 712 peaks left. Nearly all of the Elba2 sites are lost in *elba1* mutant, with 48 remained. We confirmed that loss of binding is not due to the absence of the ELBA proteins in the non-congnate *ELBA* mutant conditions, as Elba1 is normally expressed in the *elba3* mutant and Elba3 is normal in the *elba1* mutant (**Supplemental Fig. S1A**). Insv binding was not affected by any of the *elba* mutations, or vice versa (Fig. 2A**, Supplemental Fig. S2A-B**).

**Figure 2:**
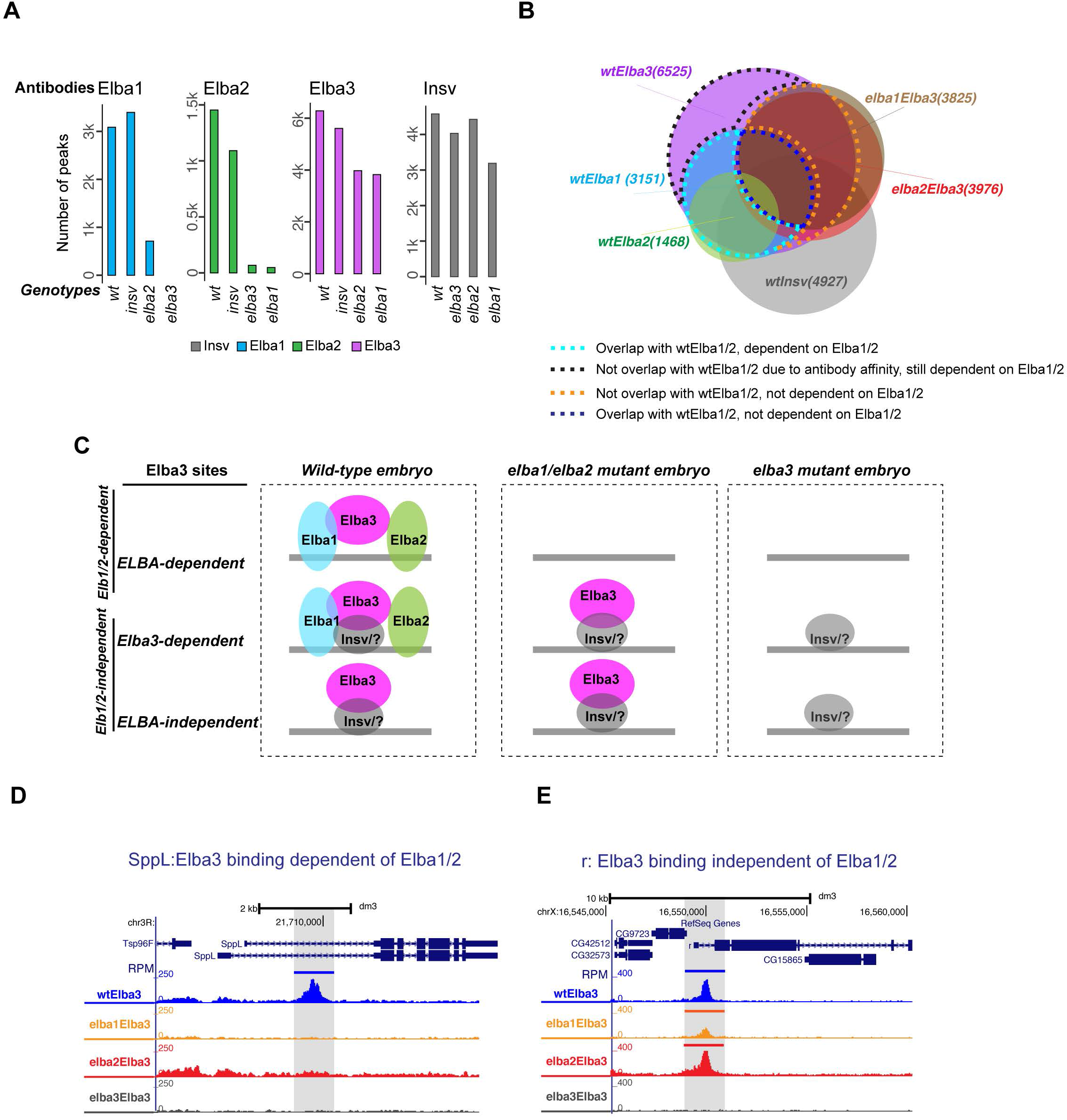
Elba1 and Elba3 binding sites are partially independent of the Elba trimeric complex. (**A**) The ChIP peaks of the *wt* or non-cognate mutants for each factor were called by using the corresponding ChIP reads against the ChIP reads in its cognate mutant. (**B**) The overlapping analysis of the Elba3 peaks in the three conditions: *wt*, *elba1* and *elba2* mutants. Four fractions of wtElba3 are highlighted with four colored dashed line according to overlapping patterns. (**C**) Illustration of three contexts where Elba3 locates in the genome. (**D**) An example locus, *SppL*, of Elba3 binding dependent of Elba1/2. (**E**) An example locus, *r*, of Elba3 binding independent of Elba1/2.

Unexpectedly, Elba3 maintains more than half of its peaks in the *elba1* and the *elba2* mutants (Fig. 2A), demonstrating that Elba3 binding to these sites does not rely on Elba1 and Elba2. To check whether these sites were bound by the ELBA complex in *wt* embryos (wtElba3), we did overlapping analysis for the peaks of each factor from *wt* condition and those for Elba3 and Elba1 from the *elba1* and the *elba2* mutants (Fig. 2B). The Elba3 sites that remained in *elba1* or *elba2* (for simplicity, referred to as *elba1/2*) mutant are mostly covered by the wtElba3 sites, suggesting that there is no global shift of Elba3 binding to new genomic locations in the absence of Elba1 and Elba2. According to the overlapping pattern, we further divided the wtElba3 sites into four fractions (highlighted by four colored dashed lines in Fig. 2B). One fraction contains the sites that disappeared in *elba1/2* mutant but do not overlap with wtElba1/2 peaks. This is presumably due to less efficient pulldown by the Elba1 and Elba2 antibodies. The second fraction covers the sites that disappeared in *elba1/2* mutants and also overlap with wtElba1/2 peaks. The third fraction contains the sites that overlap with wtElba1/2 but did not disappear in *elba1/2* mutants, and the last fraction has the sites that do not overlap with wtElba1/2 and did not disappear in *elba1/2* mutants. Based on these observations, we conclude Elba3 can bind the genome in three ways, through the ELBA complex, through protein-protein interaction with another DNA binding factor but within the ELBA complex and through protein-protein interaction with another factor without the presence of Elba1and Elba2 (Fig. 2C).

We assigned the Elba3 peaks that remained in *elba1* or *elba2* mutant as Elba1/2- independent sites (3806) and the peaks that disappeared in *elba1* or *elba2* mutant as Elba1/2-dependent (2478), exemplified by the *Sppl* and the *r* gene loci respectively (Fig. 2D-E). The Elba1/2-dependent ones are more enriched in introns, exons and distal regions and have higher frequency of motif occurrence, whereas the Elba1/2-independent sites are mostly at promoter-TSS proximal regions and contain small fraction with the motifs (**Supplemental Fig. S2C-E**). Given that Elba3 does not harbor a DNA binding domain, we asked whether Insv mediates Elba3 binding to the genome in the absence of Elba1 and Elba2. Although the fraction of overlapping peaks with Insv is higher in the independent sites than the dependent ones, half of the Elba1/Elba2-independent peaks do not overlap with Insv peaks (**Supplemental Fig. S2D**). This suggests that Insv may contribute to or enhance Elba3 binding in some but not all loci. Moreover, the Elba1/2 independent sites have higher peak scores (**Supplementary Fig. S2E**), indicating stronger binding of Elba3 to these sites. Thus, there are intrinsic differences between these two groups of Elba3 sites.

We next examined the 712 Elba1 peaks that remained in the *elba2* mutant (**Supplemental Fig. S2F-H**). This fraction of Elba1 peaks has less frequent occurrence of the Insv/Elba motifs and are more enriched in promoter-proximal regions (**Supplemental Fig. S2F**). The peak scores of these are comparable to those of the Elba1 peaks in *wt* embryos (**Supplemental Fig. S2H**), suggesting that the signals are not background noises. Interestingly, among these peaks, 496 do not overlap with the Elba1 peaks in *wt* but with the Elba3 peaks in *elba2* mutant (**Supplemental Fig. S2F**), suggesting that Elba1 may have shifted or enhanced its binding to these new loci with the help of Elba3 in the absence of Elba2. Compared to its binding sites in *wt* embryos, the Elba1 sites in *elba2* mutant are more similar to the Elba1/2-independent Elba3 sites, with a higher enrichment in promoter-TSS proximal region and fewer peaks that contain the motifs (**Supplemental Fig. S2F**). The fraction of Elba1 peaks in *elba2* mutant overlaps less with Insv, arguing that Elba1, Insv and Elba3 unlikely form a complex at target sites in the absence of Elba2 (**Supplemental Fig. S2G**).

Together, the ChIP-seq analyses revealed unexpected *in vivo* binding capacity of the three ELBA factors to the genome, in which Elba3 is essential for the complex and has the ability of targeting its genomic target sites without Elba1 and Elba2.

### ChIP-Nexus differentiates heterotrimeric binding versus homodimer binding

We reported that all the three BEN-solo proteins could bind to the symmetric site as homodimers when overexpressed in culture cells while the ELBA complex has higher affinity to the asymmetric site (Aoki et al. 2012; Dai et al. 2015; Fedotova et al. 2018). Our ChIP-seq data suggested that ELBA and Insv associate with both types of sites in the genome (**Fig.1C-D**) and that ELBA and Insv extensively overlap (Fig. 1). To overcome the issue that relatively broad ChIP-seq peaks can limit the resolution of distinguishing closely spaced factors, we performed ChIP-nexus (He et al. 2015) to better discriminate their binding preference. We used the same sets of antibodies and the same stage of *wt* embryos. As ChIP-nexus datasets lack negative control, we manually spotted coverage intensity and set stringent cut-off (FDR < 1E-10 for Elba1, Elba3 and Insv, and FDR <1E-5 for Elba2) according to signal versus background ratio. After applying this cut-off, we compared motif occurrence frequencies between ChIP-seq and ChIP-nexus. The occurrence of the Insv/ELBA motif is more frequent in the overlapping peaks of the two datasets (**Supplemental Fig. 3A**). Therefore, to ensure signal specificity of the ChIP-nexus peaks, we assigned the overlapping peaks as ChIP-nexus peaks and performed subsequent analysis on them. Compared with the ChIP-seq data alone, the ChIP-nexus peaks also have more centered distribution for both types of consensus sequences (Fig. 3A). This confirms higher specificity and resolution achieved by ChIP-nexus.

**Figure 3:**
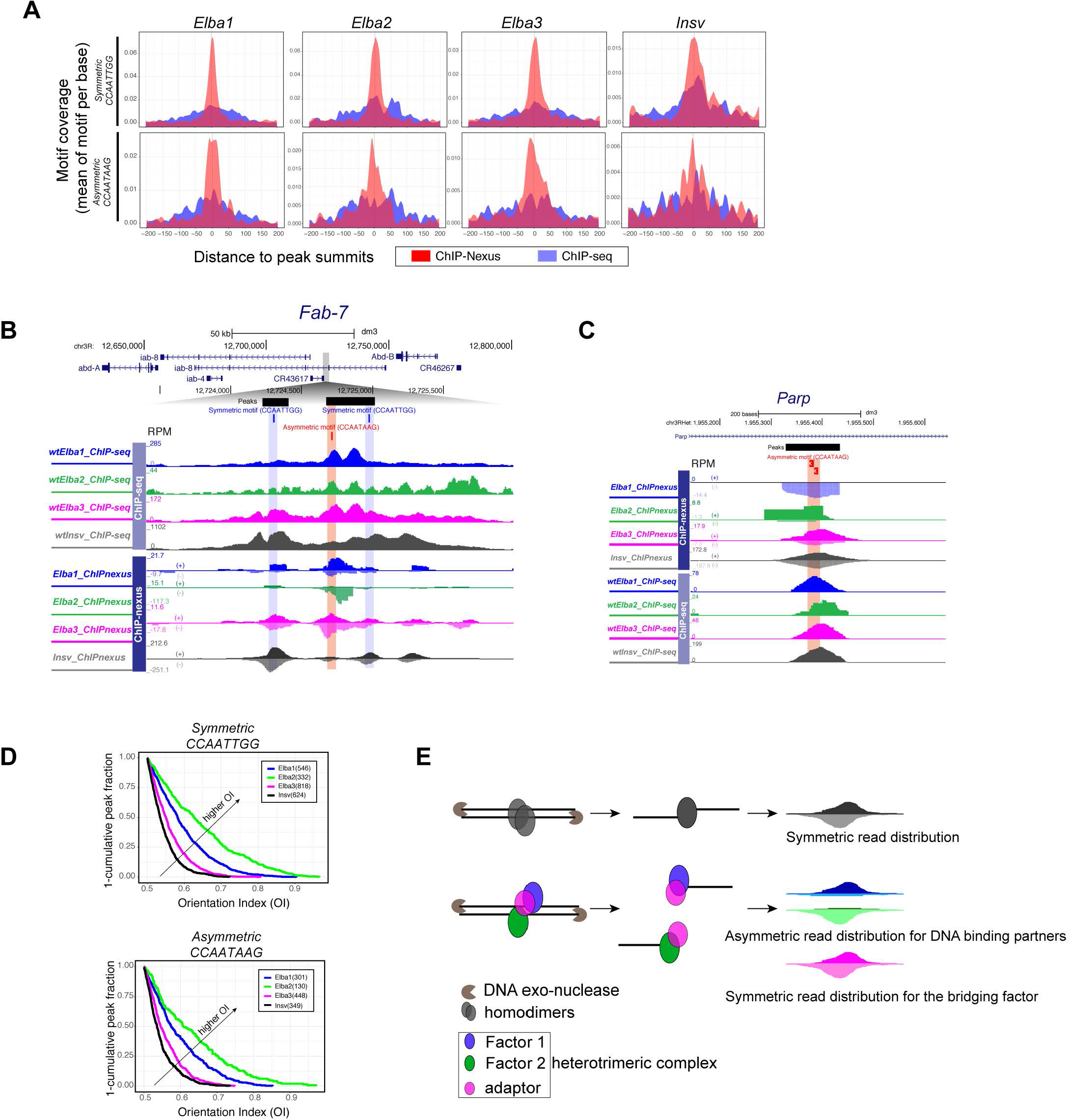
ChIP-nexus distinguishes symmetric and asymmetric TF binding sites. (**A**) Comparison of ChIP-seq and ChIP-nexus shows that ChIP-nexus has a higher frequency of motif occurrence and more centered motif distribution around the peak summits. The motif occurrence is centered at the peak summits (x-axis) and mean motif coverage in motif per base is on y-axis. (**B**) The screenshot of the Fab-7 region shows broad ChIP-seq peaks and sharp ChIP-nexus peaks. Note: asymmetric binding of Elba1 and Elba2 versus symmetric binding of Elba3 and Insv. Elba1 prefers the + strand and Elba2 the – strand of the CCAATAAG motif. (**C**) Another example locus, Parp, exhibits a similar biased strand asymmetry (OI near 1) for Elba1/2. (**D**) The orientation indexes (OI) for the peaks with the symmetric or asymmetric motifs, ranging from 0.5-1, were calculated from the ChIP-nexus reads of the four factors (see Method). The distribution of orientation indexes (OI) shows that Elba1/2 display a higher OI tendency than Elba3 and Insv. (**E**) Illustration of how ChIP-nexus can capture symmetric versus asymmetric binding patterns by homodimers versus hetero-trimeric complex.

It was shown that ELBA and Insv binding to Fab-7 contributes to the insulator function of Fab-7 (Aoki et al. 2012; Fedotova et al. 2018). The Fab-7 insulator covers one Insv/ELBA asymmetric and two symmetric sites. While the ELBA complex can bind to all three sites *in vitro* (Aoki et al. 2012; Fedotova et al. 2018), Insv binds the symmetric site more strongly (Dai et al. 2015) and the asymmetric site with lower efficiency (Fedotova et al. 2018). The ChIP-seq peaks at this locus are broad, making it difficult to differentiate ELBA or Insv specific bindings (Fig. 3B). In contrast, the ChIP-nexus peaks are sharp and show one high peak of Elba1 and Elba2 at the ELBA site, confirming that the ELBA site mediates efficient binding for Elba1 and Elba2 *in vivo*. Intriguingly, Elba1 and Elba2 peaks display strand asymmetry: Elba1 signal primarily covers the “+” strand while Elba2 signal covers the “−” strand of the CCAATAAG sequence. In contrast, Elba3 signal is relatively symmetric. Insv shows a weak peak at the ELBA site but has much stronger signal at the upstream symmetric site with no strand preference in Fab-7 (Fig. 3B). Many other individual Elba1/2-bound loci show a similar pattern, as exemplified by the *Parp1* gene: Elba1 and Elba2 preferentially bind to the “+” and the “−” strands of the two tandem asymmetric motifs respectively (Fig. 3C).

These observations prompted us to examine binding symmetry at a global level. To this end, we calculated the orientation index (OI) for every ChIP-nexus peak. OI value is determined as the ratio of the number of reads from the dominant strand to the total number of reads from both strands. Thus, OI value closer to 0.5 points to symmetric binding while closer to 1.0 indicates asymmetric binding. Similar to what we observed in individual loci, Elba3 and Insv bindings are symmetric at a global level as their OIs are mostly close to 0.5 (Fig. 3D). In contrast, the distribution of OIs of Elba1 and Elba2 are biased toward 1.0 (Fig. 3D) on both types of DNA motifs. Thus, this result suggests that the ChIP-nexus assay was able to distinguish heterotrimeric binding from homo-dimer binding (illustration in Fig. 3E).

As ChIP-nexus could improve detection of direct binding of ELBA and Insv to their cognate DNA sites, we wondered whether directly bound regions have different overlapping fractions between the four factors. We performed overlapping analysis based on the ChIP-nexus data and found that the overlapping fractions look similar to those obtained from the ChIP-seq data (**Supplemental Fig. 3B, compared with** Fig. 1B). This result confirms the presence of Insv unique-binding loci. The ChIP-nexus data also confirms that the ELBA complex can bind to the symmetric site in regions lack of Insv (**Supplemental Fig. 3C-D**) and that Insv can associate with the asymmetric site in its uniquely-bound regions (**Supplemental Fig. 3C, E**).

Within the Elba3 ChIP-nexus peaks, we identified 724 Elba1/2-dependent and 1314 Elba1/2-independent peaks. Their genomic distributions appear similar to those from ChIP-seq peaks, but the frequency of motif occurrence was substantially increased (**Supplemental Fig. 3F**). Notably, the Elba1/2-independent ChIP-nexus peaks overlap more with the Insv ChIP-nexus peaks (**Supplemental Fig. 3G**, 76%, compared with 50% in **Supplemental Fig. S2D**). This result indicates that Elba3 associates with Insv more often in embryos that lack Elba1 and Elba2, and thus provides evidence that Insv is involved in recruiting Elba3.

### All three ELBA factors repress target gene expression in *Drosophila* embryo

We previously reported that Insv represses neural genes in *Drosophila* embryos and that Elba1, Elba2 and the ELBA complex can all repress reporter gene expression in culture cells (Dai et al. 2013b; Dai et al. 2015). We asked whether Elba3 can repress transcription independent of Elba1 and Elba2. To address this, we tethered Elba3 with the Tet repressor DNA-binding domain (TetR-Elba3) and examined its activity in *Drosophila* S2 cells on a luciferase reporter driven by an actin enhancer and the *tet* Operator sites. Remarkably, Elba3 represses reporter expression with similar efficiency as the other three proteins (Fig. 4A). Given that S2 cells lack Elba1, Elba2 and Insv, this result suggests that Elba3 is able to repress transcription when brought to target promoter by another mean than by Elba1 and Elba2.

**Figure 4:**
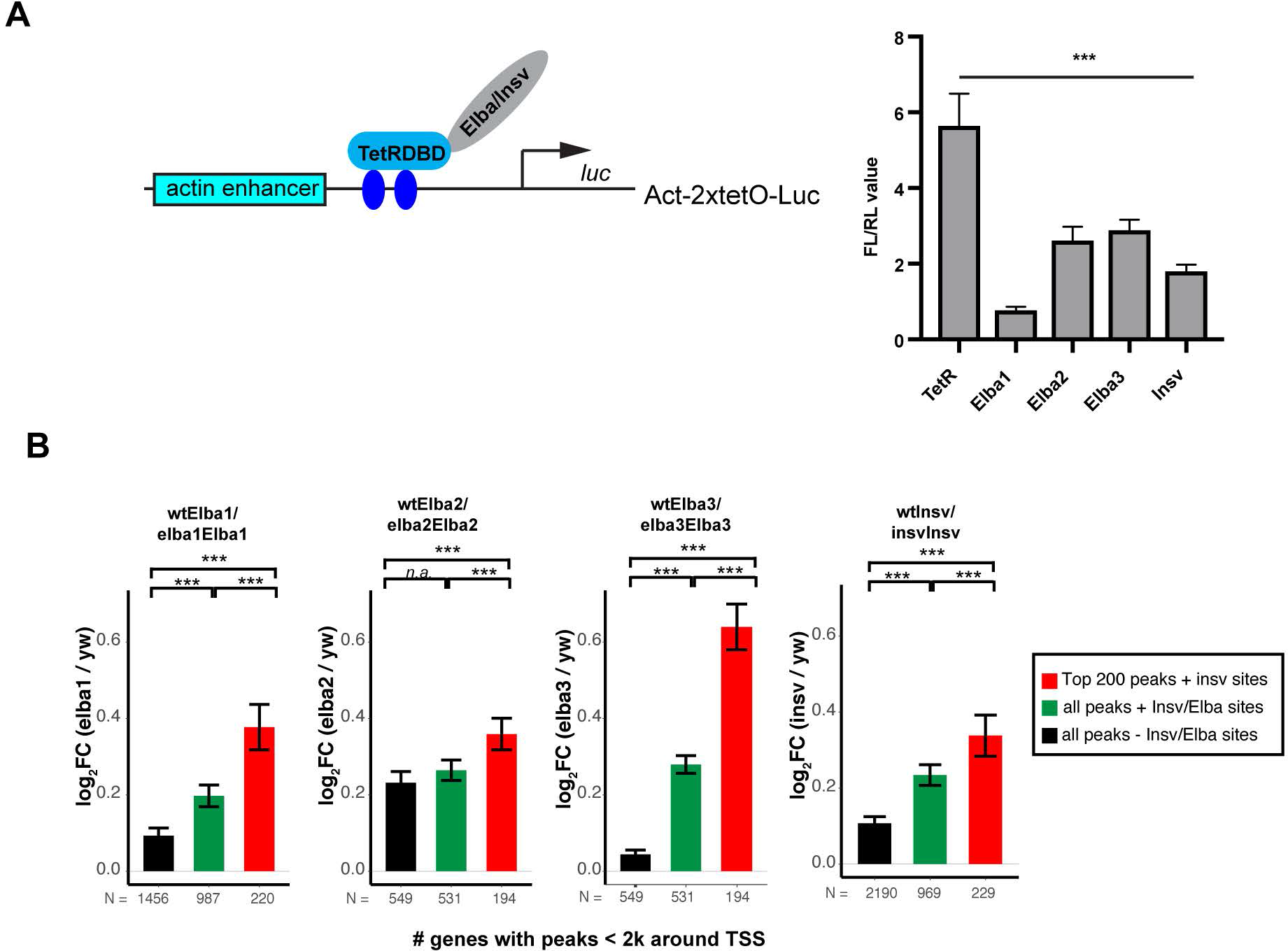
The Elba factors and Insv all repress target gene expression *in vivo*. **(A**) Luciferase reporter assays with TetR-DNA binding domain (DBD) fusion proteins. The fused Elba or Insv factors were brought to the 2xTet Operator sites by TetR-DBD and repressed reporter expression compared to the TetR-DBD alone. (B) The bar plots with Log2FC (fold change) of mutant versus wt show top bound target genes and the genes associated with the DNA consensus were more up-regulated than the other genes.

To investigate how ELBA regulates gene expression *in vivo*, we performed RNA-seq analysis to determine gene expression changes in 2-4 hour embryos between *wt* and the four mutant genotypes. Using a gene set enrichment testing (see Methods) for all the targets identified from the ELBA/Insv ChIP-seq peaks as a set, we found that the ELBA/Insv targets associated with the top200 peaks and the peaks with the Insv/ELBA motifs have a significant trend of de-repression in mutants (FDR<1E-5 for Elba1/2/3 and FDR<0.01 for Insv) but not significant in the peaks without the motifs **(Fig. 4B)**. This result demonstrates that these targets are normally repressed by ELBA/Insv in early embryo.

We then compared the genes that changed expression in these different mutants (FDR<0.2 and FC>1.3-fold). Most of the up-regulated genes in *elba2* mutant are consistently up-regulated in *elba1* and *elba3* mutants (**Supplemental Fig. S4A**). The up-regulated genes in *elba1* mutant also became up-regulated in *elba3* mutant. The overlapping pattern resembles their ChIP peak overlapping pattern (Fig. 1B), suggesting Elba3 binds to and regulates more target genes than Elba1 and Elba2. The up-regulated genes in *insv* mutant partially overlap with those in *ELBA* mutants, suggesting Insv and ELBA regulate a subset of common targets. Notably, the down-regulated genes are fewer in all the mutants and show more random overlapping (**Supplemental Fig. S4B)**, indicating down-regulation is an indirect effect, consistent with the conclusion that ELBA and Insv repress transcription.

### ELBA is required for *Drosophila* embryonic patterning

Next we sought to identify co-factors that work together with ELBA and Insv. We noticed that the highly enriched motifs in the ELBA or Insv ChIP peaks include the known binding sites for three insulator proteins, CP190, BEAF-32 and GAF (Fig. 5A). We performed pair-wise comparison for the Insv and the ELBA ChIP-seq peaks with the ChIP-ChIP peaks of CP190, BEAF-32, CTCF, GAF, Mod(Mdg4) and Su(Hw) (modEncode datasets). Consistent with our previous analysis (Dai et al. 2015), Insv co-bind with CP190, BEAF-32, CTCF and Mod(Mdg4), and shows the least overlapping with GAF and Su(Hw). The ELBA factors display similar co-occupancy patterns (Fig. 5B), suggesting that they all mainly associate with class I insulators.

**Figure 5:**
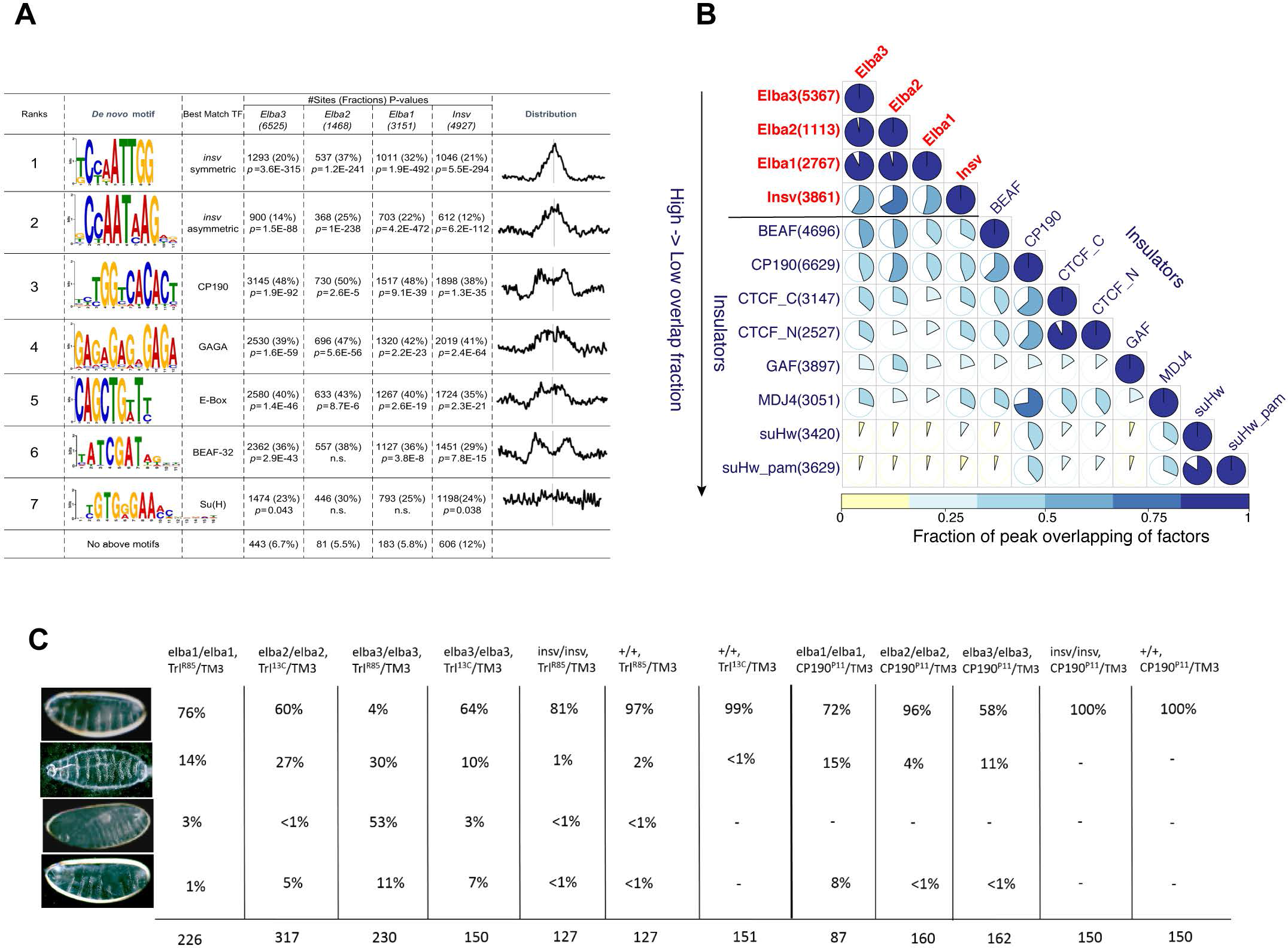
Interaction of ELBA and Insv with other insulator proteins. (**A**) The *de-novo* motif discovery analysis from the Elba and Insv peaks identified the Insv/Elba symmetric and asymmetric motifs, motifs for CP190, GAF and BEAF-32, as well as E-box and the Su(H) binding site. (**B**) The pair-wise peak overlapping matrix summarizes the genomic co-occupancy of the Elba factors, Insv and the six other insulator proteins (see Methods). Among these insulator proteins, CP190 exhibits the highest overlap with Elba and Insv, followed by BEAF-32 and CTCF. Elba and Insv have the least overlapping with GAF, mod(Mdg4), and Su(Hw). (**C**) Cuticle preps from genetic interaction tests of Elba or Insv with GAF or CP190. Animals that lack *elba* and one copy of *GAF* or *CP190* display severe patterning defects. Quantification of the normal and phenotypic embryos is shown on the right.

Knockdown of ELBA in the early embryo was shown to influence boundary activity of the HS1 element in the Fab-7 insulator (Aoki et al. 2012). Loss of ELBA or Insv also influence gene expression (Fig. 4). However, the *ELBA* and *insv* mutants are viable and do not display obvious morphologic defects. We reasoned that this could be due to redundancy with other insulator factors as they co-occupy similar genomic locations. To test this, we set genetic interaction assays between *ELBA* or *insv* and *GAF* or *CP190*. We crossed a null allele of *GAF*, *Trl*^*R85*^ (Bhat et al. 1996), a hypomorphic allele of *GAF*, *Trl*^*13C*^ (Farkas et al. 1994), and a null allele of *CP190*, *CP190*^P11^ (Pai et al. 2004), into the background of *elba1*, *elba2*, *elba3* or *insv* homozygous background, and scored for synthetic adult lethality (**Supplemental Fig. S5A**) and defects in embryonic patterning (Fig. 5C). It was shown that *insv* genetically interacts with *GAF* in the function of Fab-7 (Fedotova et al. 2018) and the Insv protein physically interacts with CP190 (Dai et al. 2015; Fedotova et al. 2019). However, we did not observe genetic interactions between *insv* and *GAF* or *CP190* in viability and early embryonic patterning. It is possible that Insv and these two factors work together in other developmental cortex. In contrast, animals homozygous for *elba3* or *elba2* in combination with heterozygous *Trl*^*R85*^ cannot survive to adulthood. Importantly, the lethality of *elba2* homozygous with *Trl*^*R85*^ is fully rescued by a pBAC transgene expressing endogenous level of *elba2* (**Supplemental Fig. S5A)**. In the combinations with heterozygous *CP190*^*P11*^, it is the homozygous *elba1* and *elba3* mutants that are lethal, suggesting distinct involvement of the three ELBA subunits with other insulator proteins in developmental processes.

A fraction of embryos mutant for *ELBA* and *Trl*^*R85*^ or *Trl*^*13C*^ also displayed severe embryonic patterning defects such as disrupted denticles and head involution, with the *elba3* and *Trl*^*R85*^ combination showing the strongest effect (only 4% normal looking embryos) (Fig. 5C). Embryos of the *elba2* and *Trl*^*R85*^ combination did not show patterning defect, presumably due to maternal contribution from *elba2* heterozygous mothers. Indeed, when embryos were produced from homozygous *elba2* and heterozygous *Trl*^*13C*^ females, a fraction of them showed patterning defects. *elba3* mutant also shows the strongest interaction with *CP190*^*P11*^, despite overall milder severity than that with *Trl* (Fig. 5C,). Importantly, *ELBA*, *Trl* or *CP190* mutant alone did not show similar defect, suggesting the interaction is specific between *ELBA* and *Trl* or *CP190*.

Together, we conclude that ELBA and Insv associate with a subset of known insulator proteins, but the Elba factors seem to be selectively needed in early embryonic development in collaboration with other insulator proteins. Importantly, even though ELBA and Insv are viable and do not exhibit substantial embryonic patterning defects, the dose-sensitive interactions we observe with other insulator proteins supports the notion that they have endogenous impacts on developmental gene regulation.

### ELBA insulates adjacent transcription units

It was suggested that Class I insulators that are enriched in gene dense regions and proximal to promoters may partition closely spaced transcription units (Negre et al. 2010). The observation that Insv and ELBA bind to this class of insulators and gene dense regions prompted us to investigate the causal role of these factors in regulating densely spaced promoters. To this end, we performed PRO-seq assay from 2-4 hr *wt* and mutant embryos and identified real-time transcripts produced by RNA PolII. We then made *de novo* PRO-seq peak calling to define actively transcribed genes in all genotypes and determined differential expression between every pair of the two adjacent promoters flanked with an ELBA or Insv ChIP peak by using the promoter reads. Notably, in *ELBA* mutant compared with wt (*yw*), there is a global reduction of expression difference between adjacent promoters (p-values adjusted by the Bonferroni correction < 0.001, **Supplemental Fig. S6A-B**). To test whether this global reduction is above background, we performed a Monte-Carlo simulation of expression difference between randomly-chosen adjacent promoters in the genome (see Method). This confirmed that the fold change between ELBA-flanked adjacent promoters is significantly higher than random. We reasoned that if the expression levels of two adjacent promoters differ more, there might be a higher need of insulation between them. This was indeed the case. For promoter pairs that differ more than 4 folds in their expression, the reduction of expression difference became even more apparent with p-values adjusted by the Bonferroni correction < 0.0001 (Fig. 6A**, Supplemental Fig. S6C)**. In contrast, for the promoter pairs that differ less than 4 folds in expression, no significant change was detected (Fig. 6B). All three types of promoter-pair configuration, convergent, tandem and divergent, showed similar trend (Fig. 6). The trend of reduction is consistent when gene-body reads were used to call differential expression (**Supplemental Fig. 6SD-E**).

**Figure 6:**
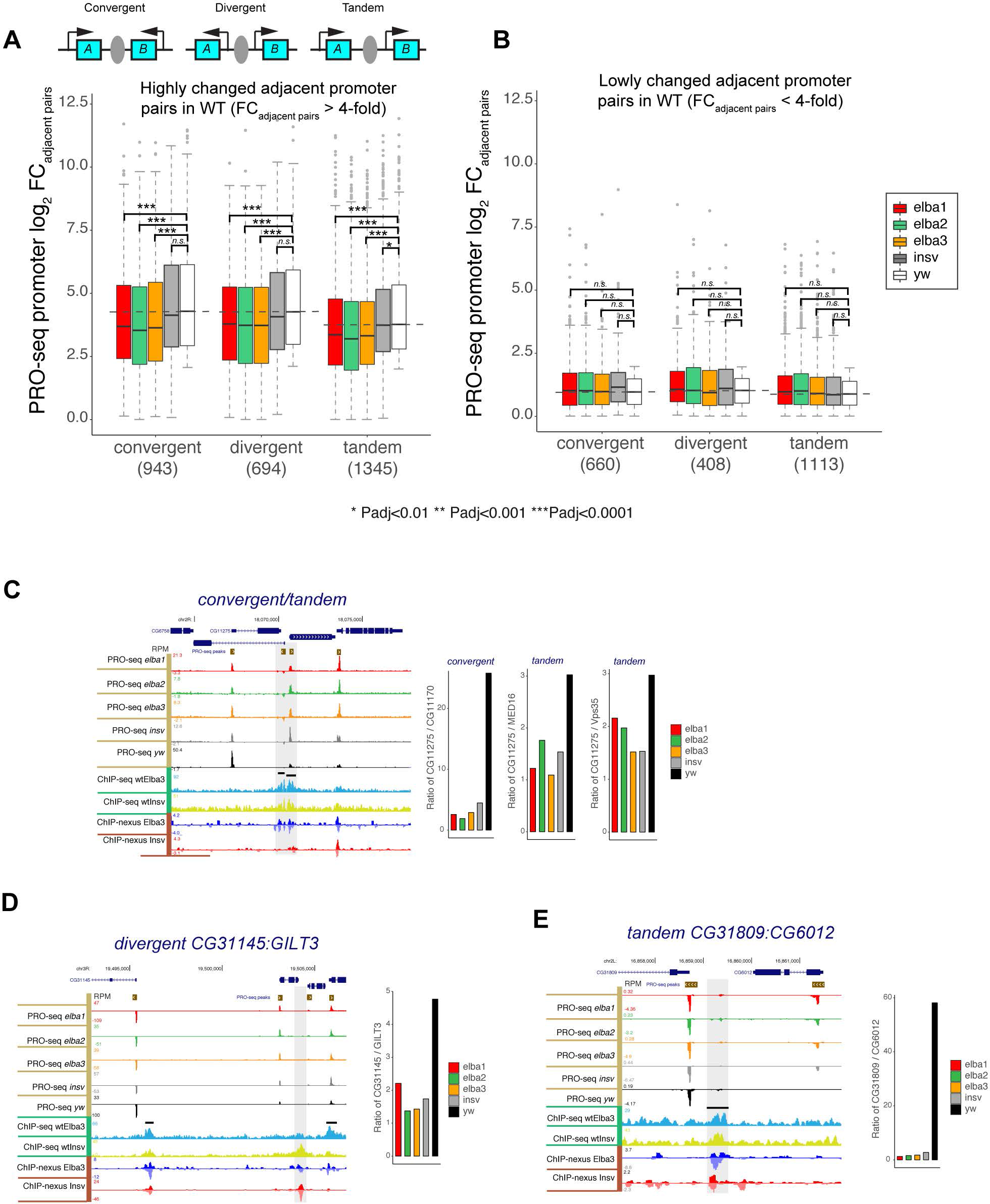
The Elba factors insulate adjacent transcription units. The PRO-seq data from *wt* and mutant embryos were used to identify real-time transcripts produced by RNA PolII in each genotype, and *de novo* PRO-seq peak call was made to define active promoters. (**A-B**) We considered three types of adjacent promoter pair configurations, convergent, divergent, and tandem, flanking an Elba/Insv peak. Differential expression between adjacent promoter pairs (absolute FC _adjacent pairs_) were divided into (**A**) the highly differentially expressed (> 4-fold) and (**B**) the lowly differentially expressed (< 4-fold) pairs in the wild-type (*wt*). **(A)** Significant reduction of expression difference between highly differentially expressed neighbor promoters in the three *elba* mutants but not in *insv* mutant. **(B)** Less differentially expressed gene pairs do not show significant changes in any of the *elba* and *insv* mutants. **(C-E)** Three example loci with convergent, divergent or tandem gene pairs flanked by the Elba/Insv binding peaks, showing the adjacent promoters became more equally expressed in *elba* and *insv* mutants compared to wt. The bar plots on the right show the ratio of expression of the two adjacent promoters in each genotype determined by PRO-seq read coverage.

*insv* mutant did not show such a global effect. However, in many individual loci, we observed a similar reduction of expression difference between Insv-bound neighbour promoters in *insv* mutant (Fig. 6C-E), suggesting that the insulation function of Insv-bound sites is present in early embryos but may be more restricted to certain gene pairs. Thus, we conclude that the ELBA factors insulate transcription units to ensure proper gene expression in *Drosophila* embryos.

### ELBA-bound elements block enhancer-promoter interaction

We reasoned if ELBA/Insv binding separates unrelated promoter-enhancer interaction in the early embryos, their bound elements may have the capacity to block enhancer interactions in ectopic setting. We sought to test this possibility by using a reporter transgene where the *LacZ* and *white* genes are controlled by both the *2xPE* and *iab-5* enhancers (Fig. 7A, (Zhou et al. 1996)). *2xPE* is an enhancer from the *twist* gene locus that will drive expression of the reporters in the ventral strip of the early embryo. iab-5 is a cis-element controlling expression of *Abd-B* in the posterior segments of the embryo. Thus, this transgenic reporter will show expression of *LacZ* and *white* in both *2xPE* and *iab-5* domains if the inserted fragment, for example the uMar spacer, does not have insulation activity (Fig. 7A). We selected eleven ELBA and Insv bound genomic loci and three control loci that do not have ELBA or Insv binding, and tested their activity in enhancer blocking (**Supplemental Fig. S7**). Two of the three control regions did not show insulation activity. The third transgene that contains the *Dpr8* region gave inconsistent results between two independent lines. In contrast, six of the bound regions show strong blocking activity on the *2xPE* enhancer from the *lacZ* gene, and weaker activity on the *iab-5* enhancer from the *white* gene (**Supplemental Fig. S7,** Fig. 7A). Therefore, many of the ELBA/Insv bound loci are insulator elements.

**Figure 7:**
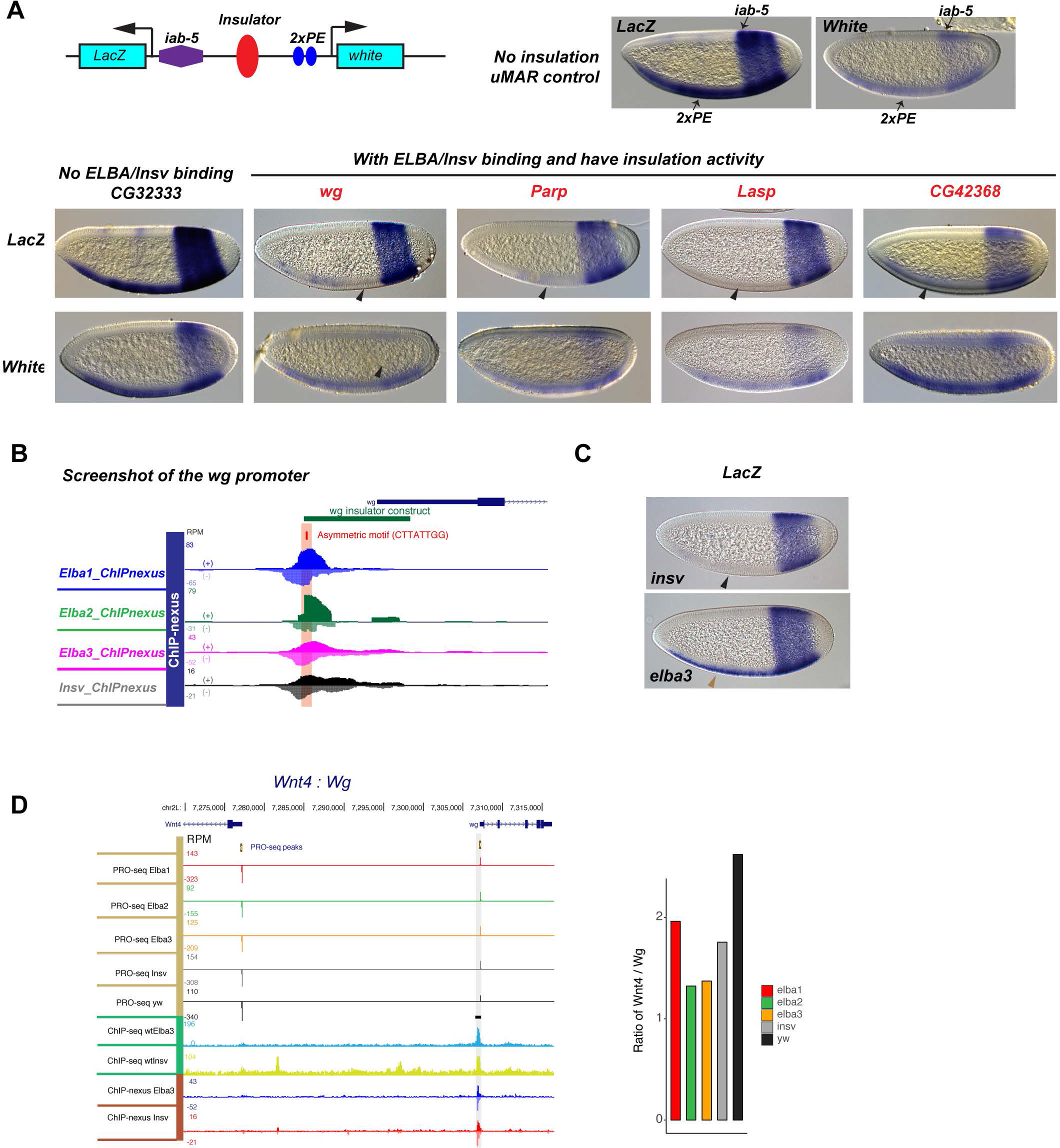
The ELBA binding site between the Wnt4 and wg divergent pair blocks enhancer-promoter interactions. **(A)** Transgenic insulator assay. *In situ* hybridization images show expression of the *lacZ* and *white* genes is driven by the 2xPE and iab-5 enhancers in the ventral and the posterior strips respectively. The insertion of a spacer sequence (uMAR) does not affect the reporter expression, neither does the CG32333 fragment that has no Elba or Insv binding. The fragments from the *wg*, *Parp* and *Lasp* loci, highlighted in red, show blocking activity, evidenced by lack of the ventral stripe of 2xPE domain in the *LacZ* staining. The iab-5 enhancer was less affected, with a weaker but visible posterior strip in *white* staining. Black arrowheads indicate the weakened or absent staining due to insulation activity of the inserted fragments. **(B**) A screenshot of the insulator in the *wg* locus, showing Elba and Insv binding and an asymmetric Insv/Elba motif. (**C**) The lacZ *in situ* staining shows loss of insulation activity of the *wg* element in the *elba3* but not *insv* mutant embryos. The black arrowhead indicates the absence of the ventral strip, showing no change in the insulation activity in the *insv* mutant. The brown arrowhead indicates recovery of the ventral strip due to loss of insulation in the *elba3* mutant. (**D**) A screenshot of the divergent pair *Wnt4* and *wg* with an Elba/Insv peak proximal to the *wg* promoter. The ratio of PRO-seq promoter expression of *Wnt* versus *wg* decreased in the *elba* and *insv* mutants compared to the *wt*.

To test whether ELBA or Insv is required for the insulation function in the reporter assay, we focused our analysis on the element in the *wg* locus that gives strongest blocking activity. This fragment contains a ELBA-type of asymmetric motif where Elba2 shows preferential binding to the strand of CTTATTGG, similar to its preference to the ELBA site in Fab-7 (Fig. 7B). The expression of the lacZ reporter remained as the same as in the wt (Fig. 7C). Remarkably, in the *elba3* mutant, the *lacZ* staining of the *2xPE*-controlled ventral strip is fully recovered, suggesting that Elba3, but not Insv, is necessary for the insulation activity of this element.

Interestingly, the *wg* promoter positions back-to-back (divergent) with the neighbour gene *Wnt4*. The ratio of expression of *Wnt4* versus *wg* decreased substantially in all the *ELBA* and *insv* mutants compared to *wt* (Fig. 7D), suggesting ELBA, probably also Insv, are required for the separation of these two genes in the endogenous context.

## Discussion

The BEN-domain containing proteins are conserved throughout metazoan, but our knowledge on the molecular and biological functions of this family is relatively poor. Here we used *Drosophila* as an *in vivo* model and investigated in depth the genomic function of the BEN-solo proteins in early embryonic development. We show that both ELBA and Insv repress transcription of direct target genes. However, only the ELBA factors play a role in early embryonic patterning together with other insulators. At a genome-wide level, ELBA is required for separating transcription of differentially-expressed neighbour genes.

### Genomic binding properties of the Drosophila BEN-solo factors

The BEN domains of Elba1, Elba2 and Insv share similar amino acid sequences and identical protein-DNA interaction sites (Dai et al. 2013b; Dai et al. 2015). However, their DNA binding activities seem to be complex. When ectopically expressed in cultured cells, all of these factors display high affinity binding to the palindromic site while only the ELBA complex is able to bind the asymmetric site (Dai et al. 2013b; Dai et al. 2015). *In vitro* translated proteins of Elba1 and Elba2 can bind to both types of motifs when additional bridging factor is present (Aoki et al. 2012; Fedotova et al. 2018). Here our ChIP-seq analyses confirm that *in vivo* Elba1 and Elba2 target the genome only through forming a heterotrimeric complex with Elba3. These results suggest that the affinity of Elba1 and Elba2 binding to DNA is weak and needs to be enhanced by additional factors. In supporting of this conclusion, our ChIP-seq analyses suggest that *in vivo* Elba3 shows strongest binding to the genome and is able to target many genomic loci including Fab-7 in the absence of Elba1 and Elba2 (Figure 2). This demonstrates that Elba3 stabilises not only the composition of the ELBA complex but also the association of ELBA with chromatin. The Elba3 protein does not have any known functional motif and not even a predictable DNA binding domain. One potential factor that can bring Elba3 to chromatin is Insv. Indeed, the Elba3 peaks that are independent of Elba1/2 overlap more with Insv peaks. But Insv should not be the only co-factor, as many of the Elba1/2 independent peaks do not overlap with Insv binding sites. Other insulator proteins with DNA binding property, such as CP190 and GAF are potential candidates that can bring Elba3 to the genome given that these factors co-occupy many genomic loci and genetic interaction.

We used high-resolution ChIP-nexus approach and confirm that Insv and ELBA factors all associate with both types of DNA motifs (Figure 3). Our ChIP-nexus analyses also provided evidence that the ELBA complex associates with DNA in an asymmetric configuration. Intriguingly, at some of the loci, such as the asymmetric sites in Fab-7 and *Parp*, Elba1 and Elba2 show + versus − strand preference. The genomic loci with asymmetric binding should represent weak association of ELBA with DNA, as strong DNA binding would allow equal pull down of the subunits with the antibody against any of the three components. There are many loci showing symmetric read distribution for Elba1 and Elba2. These sites either mediate strong binding of the complex or symmetric binding of Elba1 and Elba2 (*e.g*. as homodimers). Insv binding is always symmetric, suggesting it mostly binds to the sites as homodimers. These evidences also show ChIP-nexus can be a powerful tool to resolve binding symmetry by a heterotrimeric complex.

It will be of interest to understand how the BEN domains of all BEN proteins have evolved in DNA binding affinity and sequence specificity across species. Our previous and current work well exemplifies the approach to determine the molecular properties of a novel DNA binding protein family (Dai et al. 2013b; Dai et al. 2015).

### Functional importance of insulators in gene regulation and animal development

Activity of ELBA in the early embryo was examined by RNAi knockdown experiment where it was shown to influence early boundary activity of the HS1 element (Aoki et al. 2012). However, the effect of complete loss of ELBA in embryonic development has not been investigated. We generated the *ELBA* loss-of-function mutants and found that the *ELBA* genes are dispensable for viability. This is not surprising as other chromatin insulator proteins, such as dCTCF (Mohan et al. 2007) and BEAF-32 (Roy et al. 2007), are not required for viability. One possibility is that Drosophila utilizes multiple backup mechanisms to ensure boundary fidelity. Indeed, when one copy of *CP190* or *GAF* is removed, loss of ELBA led to drastic developmental consequences in this sensitized background (Figure 5).

Despite both ELBA and Insv associating with other known insulator proteins, such as CP190, BEAF-32 and GAF ((Dai et al. 2015), Figure 5), *ELBA* showed strong genetic interactions with CP190 and GAF in viability and early embryonic patterning while *insv* did not. This result could mean that Insv is less needed during embryonic stage and/or another unknown factor strongly compensates for its joint function with CP190 and GAF. In support of the first possibility, *insv* is required for maintaining segmentation of adult flies when the GAF sites are mutated from Fab-7 (Fedotova et al. 2018). It awaits to be identified in which other developmental contexts Insv collaborates with other insulator proteins.

Genes in the *Drosophila* genome are more compact than vertebrates. There may be a need to partition closely spaced transcription units to ensure enhancer specificity. Thanks to many years of genetic studies in *Drosophila*, a list of individual genomic loci were identified in separating enhancers or promoters (Hagstrom et al. 1996; Barges et al. 2000; Belozerov et al. 2003; Schweinsberg et al. 2004; Sultana et al. 2011; Wood et al. 2011). Insulator proteins such as GAF, CTCF, CP190 and BEAF-32 were found to mediate these activities. It was shown that the *Drosophila* insulator, BEAF-32, separates closely apposed genes with a head-to-head configuration (divergent) (Yang et al. 2012). Our results from PRO-seq analysis suggest ELBA and Insv are required to separate linked transcription units *in vivo*, evidenced by highly differentially expressed neighbour genes becoming more equally expressed in *ELBA* mutant embryos. In this case, all three types of promoter configurations, divergent, tandem and convergent, show similar requirement of ELBA. In support of the endogenous function of ELBA and Insv in blocking enhancers, a subset of genomic elements bound by ELBA and Insv are sufficient to block enhancer-promoter interaction in transgene assays (Figure 7).

In more recent years, new properties have been assigned to insulators, especially in chromatin architecture organization and long-range cis-element interactions. In this study, we focused more on the functions of ELBA and Insv in active chromatin regions because of their enrichment in close proximity to active promoters. However, we detected enrichment of ELBA and Insv in several known elements that could mediate long-range interactions, such as the homie-nhomie (Fujioka et al. 2009) and scs and scs’ loci (Kellum and Schedl 1991; Blanton et al. 2003). Future studies will be needed to determine the roles of ELBA and Insv in chromatin organization.

## Materials and Methods

### Fly strain culturing and generation of transgenes

All fly stocks were kept at 25°C. The insv mutant allele insv^23B^ was described previously (Duan et al. 2011). *elba* mutants were created using CRISPR: transgenic flies carrying single guide RNA targeting the coding sequence of each gene were crossed into the nos-Cas9 transgenic flies. Frame-shift mutations were identified by PCR and sanger sequencing. The *Trl*^*R85*^ and *Trl*^*13C*^ alleles were kindly provided by Dr. Ana Busturia (Centro de Biología Molecular “Severo Ochoa” CSIC-UAM), the *CP190*^*P11*^ allele was from Bloomington Stock Center and used for genetic interaction crosses.

For making the insulator transgenes, selected fragments were amplified and cloned into the insulator transgene backbone (kindly provided by Dr. Jumin Zhou, (Zhou et al. 1996)). The sequences of cloning oligos are provided in Supplemental table 7. All transgenic flies were created at BestGene, Inc.

### Cuticle preparation

Embryos were collected and aged to 24-36 hr before dechorionization with bleach. They were rinsed, directly mounted in 85% lactic acid, and cleared at 60°C for 3-6 hr.

### In situ hybridization

The *LacZ* and *white* probes were generated by transcription from linearized pBluescript template plasmids (kindly provided by Dr. Mattias Mannervik) with T3 or T7 RNA polymerase (Thermo Fisher) and Dig RNA labelling mix (Roche) according to manufacturer. Embryos were aged and fixed with 9% formaldehyde. in-situ hybridization was performed as previously described (Qi et al. 2008). In brief, fixated embryos were permeabilized with xylene and re-hydrated as well as post-fixated with 5% formaldehyde in PBT (1x PBS, 0.1% Tween-20) for 25 min. Embryos were treated with proteinase K (4μg/ml) for 8 min, followed by another round of post-fixation for 25 min, before hybridization with the probes at 55°C for over-night in hybridization buffer (50% formamide, 5x SSC, 100μg/ml sonicated boiled ssDNA, 0.1% Tween 20). Samples were incubated with alkaline-phosphatase-labelled anti-Digoxigenin antibody (1:2000, Roche) over night at 4°C, and developed with 0.6mg/ml Nitrotetrazolium Blue chloride (NBC) and 0.3mg/ml 5-Bromo-4-chloro-3-indolyl phosphate disodium salt (BCIP). Samples were dehydrated by repeated washes in ethanol, rinsed in xylene and mounted in Permount (Fisher).

### Cell culture and luciferase assay

To generate the TetR-DBD fusions with the Elba factors and Insv, the open reading frames of Elba1, Elba2 Elba3 and Insv were PCR amplified and cloned into the pAC-TetR vector. All transfections were performed using *Drosophila* S2-R+ cells grown in Schneider *Drosophila* medium containing 10% fetal calf serum. Cells were co-transfected with TetR fusion, 2xTetO-Firefly luciferase and pAc-Renilla plasmids in 96- well plate using the Effectene Transfection kit (Qiagene). Luciferase assays were performed and measured as previously described (Dai et al. 2013b) and using the Dual Luciferase Assay System (Promega). Expression was calculated as the ratio between the firefly and *Renilla* luciferase activities.

### ChIP-seq assay and peak calling

Chromatin immunoprecipitation (ChIP) was done mostly as previously described (Dai et al. 2013b) with one modification. In the fixation step, 2.5 mM DSG (Di(N-succinimidyl) glutarate) (Sigma) was added to the fixation buffer containing 1.8% of formaldehyde. The rest of ChIP steps were unchanged. The Elba antisera were tested in ChIP previously (Aoki et al. 2014) and kindly provided by Dr. Paul Schedl (Princeton University). For each ChIP reaction, 5ul of antibody and 50 ul of embryos were used. ChIP-seq libraries were made using the NEBNext Ultra™ II DNA Library Prep Kit.

The ChIP-seq samples were mapped to the *Drosophila melanogaster* (dm3) genome assembly using Bowtie2 with the default parameters, after the adaptor trimming by Trimmomatic. The uniquely mapped reads with a mapping quality MAPQ ≥ 20 were used for further analysis. For all ChIP-seq samples, we generated coverage tracks at 1-nt resolution and normalized to the library sizes to give read per million (RPM) in “bigwig” format. We further generated the coverage differential tracks for four factors by subtracting mutant from wt coverage (log2 wt/mutant).

For each of the four factors, the peak calling was performed by the ChIP-seq reads of wt or a mutant condition to its own mutant ChIP or IgG or Input. The peaks were called using MACS2 (Zhang et al. 2008) with default parameters and the confident peaks were determined by an FDR < 1%. The peaks overlapped with *Drosophila* blacklist were also removed. Peak overlap analysis was performed by “mergePeaks” function in Homer2 package with the default parameters.

The *de novo* motif search was performed for all the called peaks for each factor by MEME-ChIP (Machanick and Bailey 2011). We extended 500 bps of the summits of the called peaks for each factor in each direction, and run MEME-ChIP to search for 5-15 nt motifs in the central regions (100 nucleotides) using default parameters. The summits of the called peaks for each factor were extended by 500 nucleotides in each direction, and MEME-chip was run to search for 5-15 nt motifs in the central regions (100 nucleotides) using default parameters.

Pairwise comparison was done for the ChIP-seq peaks of Elba and Insv factors with the modEncode insulator datasets (Negre et al. 2010) that include ChIP-ChIP data for CP190, BEAF32, CTCT, GAF, Mod(Mdg4) and Su(Hw). As the ChIP-seq peaks are generally narrower than ChIP-chip regions, we used the summit of ChIP-ChIP regions with 100 nt extension on each side for the overlapping analysis. The range of 50 nt distance between the two peak summits was used. The overlap fraction of set1 and set2 peaks was calculated by #overlapped peaks divided by minimum of #set1 peaks and #set2 peaks.

### ChIP-nexus and analysis

ChIP-nexus was performed following the protocol step by step described previously (He et al. 2015). 20 ul of each antibody and 200 ul of embryos were used in each ChIP-nexus reaction. All the ChIP libraries were sequenced on the Illumina Hiseq2500 platform with 1×50 bp SR configuration.

Before aligning the ChIP-nexus reads to the genome, the 5’ fixed barcode (1-5) was first removed and the random 4nt barcode was retained for each read. After the 3’ adaptor trimming by Trimmomatic, the sequencing reads were collapsed to only include unique reads. The random 4nt barcode was further removed and the reads with at least 22nt were retained for mapping. We mapped the reads using bowtie with the parameter setting “-k 1 -m 1 -v 2 --best --strata”. Similar to ChIP-seq, we generated normalized coverage tracks separately for each strand in “bigwig” format. Similar to the ChIP-seq data, the ChIP-nexus peak calling was performed by MACS2 using the default parameter. To obtain highly confident binding sites for each factor, we required the binding sites to be called by both ChIP-seq and ChIP-nexus and set a highly stringent cut-off (FDR < 1E-10 for Elba1, Elba3 and Insv, and FDR <1E-5 for Elba2).

To examine asymmetry of the binding sites, we calculated an orientation index (OI) for each binding site by ChIP-nexus for each factor. OI was defined by maximum #reads between two strands divided by sum of reads of two strands, max(forward,reverse)/sum(forward,reverse), ranging from 0.5 to 1.

### RNA-seq and analysis

Total RNA was extracted from stages 2-4 hr embryos using Trizol reagent (Invitrogen). RNA quality of three biological replicates was tested by Agilent Bioanalyzer. RNA-seq libraries were made using the Illumina Truseq Total RNA library Prep Kit LT. Sequencing was performed on the Illumina Hiseq2500 platform.

After trimming the adaptor sequences using Trimmomatic, for each factor, the RNA-seq reads from the replicated wild type (x3) and mutant samples (x3) were mapped to the *Drosophila melanogaster* (dm3) genome assembly genome assembly using HISAT2. RNA-seq signal was normalized by the TMM method implemented in the Limma Bioconductor library (Ritchie et al. 2015). Gene annotation was obtained from the FlyBase dm3 gene annotation. Differentially expressed mRNAs between BEN factors mutants versus wild type were identified, and FDR (Benjamini-Hochberg) was estimated, using Limma.

To test whether a set of genes are significantly changed (up- or down-regulated as set) amongst the differentially expressed (DE) genes from wild type and mutant RNA-seq data, gene set enrichment testing function “camera” in the R limma package was used [Ritchie ME et al, 2015]. It is a ranking based gene set test accounting for inter-gene correlation, to test whether the called peaks by ChIP-seq, top200 peaks, all peaks with or without insv motifs, are significantly changed as a set.

### PRO-seq assay and analysis

The PRO-seq procedure was performed according to the previously reported method (Kwak et al. 2013). Embryos were collected from *yw*, *elba*, and *insv* mutants and aged for 3-4 hr. After the run-on reaction, Biotin-labelled RNAs were purified, enriched and cloned into cDNA libraries. In the PCR amplification step, 14 cycles were used to enrich the cDNAs for sequencing. Barcoded libraries were pooled and sequenced on the Illumina Hiseq2500 platform with 1×50 bp SR configuration.

The adaptors were first trimmed from the sequencing reads by cutadapt software and the reads with at least 15 nt were retained. We then removed reads that mapped to rRNAs and the remaining reads were further mapped to the *Drosophila melanogaster* (dm3) genome assembly using BWA with the default parameters. The PRO-seq normalized coverage tracks with separate strands were generated for each factor. To detect *de novo* transcripts from PRO-seq, we combined all genotypes and adapted the Homer2 (Heinz et al. 2010) GRO-seq transcript identification method using a parameter setting “findPeaks -style groseq -tssFold 4 -bodyFold 3”. The pausing regions (promoter region) were defined from the *de novo* transcript starts to 200nt downstream, and gene body regions were defined from 400nt downstream to the end of the *de novo* transcripts. The *de novo* transcripts having a promoter expression of greater than 1 transcript per million (TPM) were retained for further analysis.

### Analysis of ELBA/Insv factors acting as insulators

For each Elba/Insv binding site in the high confident binding set, which was called by both ChIP-seq and ChIP-nexus (see Method above), we looked for the adjacent upstream and downstream PRO-seq promoter pair and calculated absolute differential expression between them (abs log2FC adjacent pair). We classified adjacent promoter pairs flanking an Elba/Insv peak into 3 types: convergent, divergent, and tandem.

To test whether the change of differential expression between the adjacent pairs is above background, we performed a Monte-Carlo simulation. We randomly located the same number of regions with the same length as the Elba/Insv ChIP peaks in the same chromosome and repeated the random selection and calculation 2000 times. P-values were calculated by dividing the number of instances that show bigger fold change between the random adjacent genes than that between the Elba/Insv bound genes by 2000 iterations. These were done separately for convergent, divergent, and tandem pairs in each of wt and four mutants.

## Acknowledgements

We thank the imaging facility (IFSU) of Stockholm University and the National Genomic Institute of Scilife Laboratories, Sweden, for providing service and support. The Bloomington Stock Center and the *Drosophila* community provided important reagents and fly stocks. J.W. was supported by the Australian Research Council (ARC) Future Fellowship (FT60100143). The project was supported by the Young Investigator grant from Swedish Research Council (Vetenskapsrådet, 2014-5584) to Q.D.

## Author contributions

QD conceived and designed the project. EL helped the initiation of the project. MU and QD performed most of the experiments. HW performed the luciferase assays. CZ generated the constructs for luciferase assays. SK generated the *elba* mutants. PS and TA provided valuable information on the Elba antibodies. JW performed all the computational analysis. QD and JW analysed and interpreted the data with input from MU and HW. QD wrote the manuscript with help from JW and input from other co-authors.

## References

Abhiman S, Iyer LM, Aravind L. 2008. BEN: a novel domain in chromatin factors and DNA viral proteins. Bioinformatics 24: 458–461.

Aoki T, Sarkeshik A, Yates J, Schedl P. 2012. Elba, a novel developmentally regulated chromatin boundary factor is a hetero-tripartite DNA binding complex. Elife 1: e00171.

Aoki T, Wolle D, Preger-Ben Noon E, Dai Q, Lai EC, Schedl P. 2014. Bi-functional cross-linking reagents efficiently capture protein-DNA complexes in Drosophila embryos. Fly (Austin) 8: 43–51.

Barges S, Mihaly J, Galloni M, Hagstrom K, Muller M, Shanower G, Schedl P, Gyurkovics H, Karch F. 2000. The Fab-8 boundary defines the distal limit of the bithorax complex iab-7 domain and insulates iab-7 from initiation elements and a PRE in the adjacent iab-8 domain. Development 127: 779–790.

Bell AC, West AG, Felsenfeld G. 1999. The protein CTCF is required for the enhancer blocking activity of vertebrate insulators. Cell 98: 387–396.

Belozerov VE, Majumder P, Shen P, Cai HN. 2003. A novel boundary element may facilitate independent gene regulation in the Antennapedia complex of Drosophila. The EMBO journal 22: 3113–3121.

Bhat KM, Farkas G, Karch F, Gyurkovics H, Gausz J, Schedl P. 1996. The GAGA factor is required in the early Drosophila embryo not only for transcriptional regulation but also for nuclear division. Development 122: 1113–1124.

Blanton J, Gaszner M, Schedl P. 2003. Protein:protein interactions and the pairing of boundary elements in vivo. Genes Dev 17: 664–675.

Dai Q, Andreu-Agullo C, Insolera R, Wong LC, Shi SH, Lai EC. 2013a. BEND6 is a nuclear antagonist of Notch signaling during self-renewal of neural stem cells. Development 140: 1892–1902.

Dai Q, Ren A, Westholm JO, Duan H, Patel DJ, Lai EC. 2015. Common and distinct DNA-binding and regulatory activities of the BEN-solo transcription factor family. Genes Dev 29: 48–62.

Dai Q, Ren A, Westholm JO, Serganov A, Patel DJ, Lai EC. 2013b. The BEN domain is a novel sequence-specific DNA binding domain conserved in neural transcriptional repressors. Genes Dev 27: 602–614.

Dorsett D. 1993. Distance-independent inactivation of an enhancer by the suppressor of Hairy-wing DNA-binding protein of Drosophila. Genetics 134: 1135–1144.

Duan H, Dai Q, Kavaler J, Bejarano F, Medranda G, Negre N, Lai EC. 2011. Insensitive is a corepressor for Suppressor of Hairless and regulates Notch signalling during neural development. The EMBO journal 30: 3120–3133.

Farkas G, Gausz J, Galloni M, Reuter G, Gyurkovics H, Karch F. 1994. The Trithorax-like gene encodes the Drosophila GAGA factor. Nature 371: 806–808.

Fedotova A, Aoki T, Rossier M, Mishra RK, Clendinen C, Kyrchanova O, Wolle D, Bonchuk A, Maeda RK, Mutero A et al. 2018. The BEN Domain Protein Insensitive Binds to the Fab-7 Chromatin Boundary To Establish Proper Segmental Identity in Drosophila. Genetics 210: 573–585.

Fedotova A, Clendinen C, Bonchuk A, Mogila V, Aoki T, Georgiev P, Schedl P. 2019. Functional dissection of the developmentally restricted BEN domain chromatin boundary factor Insensitive. Epigenetics Chromatin 12: 2.

Fujioka M, Wu X, Jaynes JB. 2009. A chromatin insulator mediates transgene homing and very long-range enhancer-promoter communication. Development 136: 3077–3087.

Gerasimova TI, Gdula DA, Gerasimov DV, Simonova O, Corces VG. 1995. A Drosophila protein that imparts directionality on a chromatin insulator is an enhancer of position-effect variegation. Cell 82: 587–597.

Geyer PK, Corces VG. 1992. DNA position-specific repression of transcription by a Drosophila zinc finger protein. Genes Dev 6: 1865–1873.

Hagstrom K, Muller M, Schedl P. 1996. Fab-7 functions as a chromatin domain boundary to ensure proper segment specification by the Drosophila bithorax complex. Genes Dev 10: 3202–3215.

He Q, Johnston J, Zeitlinger J. 2015. ChIP-nexus enables improved detection of in vivo transcription factor binding footprints. Nat Biotechnol 33: 395–401.

Heinz S, Benner C, Spann N, Bertolino E, Lin YC, Laslo P, Cheng JX, Murre C, Singh H, Glass CK. 2010. Simple combinations of lineage-determining transcription factors prime cis-regulatory elements required for macrophage and B cell identities. Mol Cell 38: 576–589.

Holdridge C, Dorsett D. 1991. Repression of hsp70 heat shock gene transcription by the suppressor of hairy-wing protein of Drosophila melanogaster. Mol Cell Biol 11: 1894–1900.

Kaul-Ghanekar R, Jalota A, Pavithra L, Tucker P, Chattopadhyay S. 2004. SMAR1 and Cux/CDP modulate chromatin and act as negative regulators of the TCRbeta enhancer (Ebeta). Nucleic acids research 32: 4862–4875.

Kellum R, Schedl P. 1991. A position-effect assay for boundaries of higher order chromosomal domains. Cell 64: 941–950.

Kellum R, Schedl P. 1992. A group of scs elements function as domain boundaries in an enhancer-blocking assay. Mol Cell Biol 12: 2424–2431.

Korutla L, Degnan R, Wang P, Mackler SA. 2007. NAC1, a cocaine-regulated POZ/BTB protein interacts with CoREST. Journal of neurochemistry 101: 611–618.

Korutla L, Wang PJ, Mackler SA. 2005. The POZ/BTB protein NAC1 interacts with two different histone deacetylases in neuronal-like cultures. Journal of neurochemistry 94: 786–793.

Kwak H, Fuda NJ, Core LJ, Lis JT. 2013. Precise maps of RNA polymerase reveal how promoters direct initiation and pausing. Science 339: 950–953.

Kyrchanova O, Georgiev P. 2014. Chromatin insulators and long-distance interactions in Drosophila. FEBS Lett 588: 8–14.

Lutz M, Burke LJ, Barreto G, Goeman F, Greb H, Arnold R, Schultheiss H, Brehm A, Kouzarides T, Lobanenkov V et al. 2000. Transcriptional repression by the insulator protein CTCF involves histone deacetylases. Nucleic Acids Res 28: 1707–1713.

Machanick P, Bailey TL. 2011. MEME-ChIP: motif analysis of large DNA datasets. Bioinformatics 27: 1696–1697.

Mohan M, Bartkuhn M, Herold M, Philippen A, Heinl N, Bardenhagen I, Leers J, White RA, Renkawitz-Pohl R, Saumweber H et al. 2007. The Drosophila insulator proteins CTCF and CP190 link enhancer blocking to body patterning. The EMBO journal 26: 4203–4214.

Negre N, Brown CD, Shah PK, Kheradpour P, Morrison CA, Henikoff JG, Feng X, Ahmad K, Russell S, White RA et al. 2010. A comprehensive map of insulator elements for the Drosophila genome. PLoS genetics 6: e1000814.

Pai CY, Lei EP, Ghosh D, Corces VG. 2004. The centrosomal protein CP190 is a component of the gypsy chromatin insulator. Mol Cell 16: 737–748.

Perez-Juste G, Garcia-Silva S, Aranda A. 2000. An element in the region responsible for premature termination of transcription mediates repression of c-myc gene expression by thyroid hormone in neuroblastoma cells. J Biol Chem 275: 1307–1314.

Phillips-Cremins JE, Corces VG. 2013. Chromatin insulators: linking genome organization to cellular function. Mol Cell 50: 461–474.

Qi D, Bergman M, Aihara H, Nibu Y, Mannervik M. 2008. Drosophila Ebi mediates Snail-dependent transcriptional repression through HDAC3-induced histone deacetylation. The EMBO journal 27: 898–909.

Rampalli S, Pavithra L, Bhatt A, Kundu TK, Chattopadhyay S. 2005. Tumor suppressor SMAR1 mediates cyclin D1 repression by recruitment of the SIN3/histone deacetylase 1 complex. Molecular and cellular biology 25: 8415–8429.

Ritchie ME, Phipson B, Wu D, Hu Y, Law CW, Shi W, Smyth GK. 2015. limma powers differential expression analyses for RNA-sequencing and microarray studies. Nucleic Acids Res 43: e47.

Roy S, Gilbert MK, Hart CM. 2007. Characterization of BEAF mutations isolated by homologous recombination in Drosophila. Genetics 176: 801–813.

Sathyan KM, Shen Z, Tripathi V, Prasanth KV, Prasanth SG. 2011. A BEN-domain-containing protein associates with heterochromatin and represses transcription. J Cell Sci 124: 3149–3163.

Schweinsberg S, Hagstrom K, Gohl D, Schedl P, Kumar RP, Mishra R, Karch F. 2004. The enhancer-blocking activity of the Fab-7 boundary from the Drosophila bithorax complex requires GAGA-factor-binding sites. Genetics 168: 1371–1384.

Sultana H, Verma S, Mishra RK. 2011. A BEAF dependent chromatin domain boundary separates myoglianin and eyeless genes of Drosophila melanogaster. Nucleic Acids Res 39: 3543–3557.

Valenzuela L, Kamakaka RT. 2006. Chromatin insulators. Annu Rev Genet 40: 107–138.

Vazquez J, Schedl P. 2000. Deletion of an insulator element by the mutation facet-strawberry in Drosophila melanogaster. Genetics 155: 1297–1311.

Wood AM, Van Bortle K, Ramos E, Takenaka N, Rohrbaugh M, Jones BC, Jones KC, Corces VG. 2011. Regulation of chromatin organization and inducible gene expression by a Drosophila insulator. Mol Cell 44: 29–38.

Xuan C, Wang Q, Han X, Duan Y, Li L, Shi L, Wang Y, Shan L, Yao Z, Shang Y. 2013. RBB, a novel transcription repressor, represses the transcription of HDM2 oncogene. Oncogene 32: 3711–3721.

Yang J, Ramos E, Corces VG. 2012. The BEAF-32 insulator coordinates genome organization and function during the evolution of Drosophila species. Genome research 22: 2199–2207.

Zhang Y, Liu T, Meyer CA, Eeckhoute J, Johnson DS, Bernstein BE, Nusbaum C, Myers RM, Brown M, Li W et al. 2008. Model-based analysis of ChIP-Seq (MACS). Genome Biol 9: R137.

Zhou J, Barolo S, Szymanski P, Levine M. 1996. The Fab-7 element of the bithorax complex attenuates enhancer-promoter interactions in the Drosophila embryo. Genes Dev 10: 3195–3201.

