## Supplemental Figures for "BEN-solo factors partition active chromatin to ensure proper gene activation in *Drosophila*"

**Supplemental Figures 1-7**

**Figure S1:** Relevant to main Figure 1.

**Figure S2:** Relevant to main Figure 2.

**Figure S3:** Relevant to main Figure 3.

**Figure S4:** Relevant to main Figure 4.

**Figure S5:** Relevant to main Figure 5

**Figure S6:** Relevant to main Figure 6.

**Figure S7:** Relevant to main Figure 7.

**Supplemental Figure legends:**

**Figure S1:** Relevant to main Figure 1.

**(A)** Schematics of the disruption sites of the *elba* frame-shift mutations, and images of immunofluorescent staining on *wt*, *elba1* and *elba3* mutant embryos against the Elba1 and Elba3 antibodies. Nuclear staining of Elba1 is lost in *elba1* mutant but present in other genotypes. Similarly, nuclear staining of Elba3 is only lost in *elba3* mutant. DAPI is used to label the nucleus. **(B)** Normalized coverage of the ChIP-seq signal in wild-type against mutant, IgG or Input centered at TSS is shown for each of the four Elba and Insv factors. The fraction of motifs in the called peaks indicates that the wild type ChIP against mutant ChIP gave the highest motif enrichment. **(C)** Comparison of the genomic distribution of Elba3 and Insv binding sites shows that the majority of Elba3-unique sites located in the promoter-TSS proximal regions while the Insv-unique sites are more evenly distributed in different genomic regions. **(D)** Comparison of the motif enrichment of Elba3 and Insv binding regions shows that Insv-unique peaks contain a larger fraction of the know Insv/Elba motifs.

**Figure S2:** Relevant to main Figure 2.

**(A)** For each of four factors, the number of peaks was obtained by using ChIP reads from wt or mutant conditions against its own mutant ChIP. This supplement figure re-grouped the number of peaks in Figure 2 to have the same genotype in each sub-figure. The gene names with all the letters in lower case denote genotypes and the names with the first letter upper-case denote antibodies in ChIP. For each of the four factors, number of peaks was obtained by using the ChIP-seq reads of wt against its own mutant ChIP. **(B)** Insv binding in three conditions: wt, *elba2*, and *elba3* mutants show that Insv binding does not rely on Elba factors. **(C)** Comparison between Elba1/2-dependent and Elba1/2-independent Elba3-binding sites. The Elba1/2-dependent set has a higher fraction containing the Insv/Elba motifs compared to the Elba1/2-independent set. The Elba1/2-independent sites are enriched in promoter-proximal regions while the Elba1/2-dependent sites are mostly distributed in introns, exons, and distal regions. **(D)** Comparison of the Elba3 sites with the Insv sites shows that 50% Elba1/2-independent Elba3 binding sites co-bind with Insv while only 25% Elba1/2-dependent Elba3 binding sites co-bind with Insv. **(E)** Cumulative distributions of the peak scores for these two sets demonstrates that the Elba1/2-dependent set has significant lower peak scores than the Elba1/2-independent one. **(F)** Comparison of Elba1 binding in wt (wt/Elba1) and in the *elba2* mutant (elba2/Elba1). elba2/Elba1 has a smaller fraction containing the Insv/Elba motif and is more enriched in the promoter-proximal regions compared to wt/Elba1. **(G)** Overlapping analysis of Elba1 sites with Insv sites shows that the elba2/Elba1 set overlaps less with Insv than the wt/Elba1 set and the Elba3 sites in the *elba2* mutant (elba2/Elba3). **(H)** Comparison of cumulative distributions of the peak scores shows that the Elba1 sites in *elba2* mutant have comparable peak scores to the Elba1 sites in *wt* or the Elba3 sites in *elba2* mutant set.

**Figure S3:** Relevant to main Figure 3.

**(A)** For each of the four factors, the comparison of motif occurrence frequency between ChIP-seq and ChIP-nexus peaks shows that the overlapping fraction has a higher frequency of motif occurrence (purple) than the ChIP-seq-unique (green) and ChIP-nexus unique (orange) sets. **(B)** A Venn diagram shows overlapping fractions of the binding sites detected in the ChIP-nexus/ChIP-seq overlapping fraction for the four factors. **(C)** Similar to (B), but the peaks are divided into the ones with either motif or with both. **(D)** An example locus, *ERR*, shows Elba localizes at a symmetric site without Insv. **(E)** An example locus, Kirre, shows Insv localizes at an asymmetric site without Elba. **(F)**

Comparison of the Elba1/2 dependent and independent Elba3 peaks determined by both ChIP-seq and ChIP-nexus shows an increased frequency of motif occurrence when ChIP-nexus is included, compared to Supplemental Fig. S2. **(G)** Elba1/2 independent Elba3 sites also show an increased overlap fraction with Insv (from ~50% to ~75%), indicating that Insv may be required for recruiting Elba3.

**Figure S4:** Relevant to main Figure 4.

Overlapping analyses of differentially expressed genes in the four mutant genotypes compared to wild-type (FDR< 0.2 and FC >1.3-fold), for up-regulated **(A)** and down-regulated genes **(B).**

**Figure S5:** Relevant to main Figure 5.

(**A**) Lethality tests for various genetic combinations. The lethal genotypes are indicated in red.

**Figure S6:** Relevant to main Figure 6.

**(A)** A global reduction of PRO-seq expression change ($FC_{\text{adjacent pair}}$) between the adjacent promoters is detected in mutant compared to wt for the three Elba factors but not for Insv. All three types of gene pair configuration, convergent, divergent, and tandem show a similar trend. **(B)** Quantification of the reduction in (A, all Elba/Insv flanked promoters) shows the significant global deduction for the Elba factors. **(C)** Quantification of the reduction for highly differentially expressed gene pairs (expression difference > 4-fold) in Figure 6A shows stronger effects than all the active pairs in (A). **(D-E)** Similar to the promoter pairs, using RRO-seq gene body expression, the expression changes between adjacent pairs show a significant reduction in the three *elba* mutants but not in *insv* mutant for the highly differentially expressed (> 4-fold) genes. No change is detected for the lowly differentially expressed (< 4-fold) genes.

**Figure S7:** Relevant to main figure 7.

**(A)** The other transgenes in transgenic insulator assay. *In situ* hybridization images show expression of the *lacZ* and *white* genes is driven by the 2xPE and iab-5 enhancers. The reporters with detectable effects are highlighted in red. (**B**) A summary of all transgenes tested.

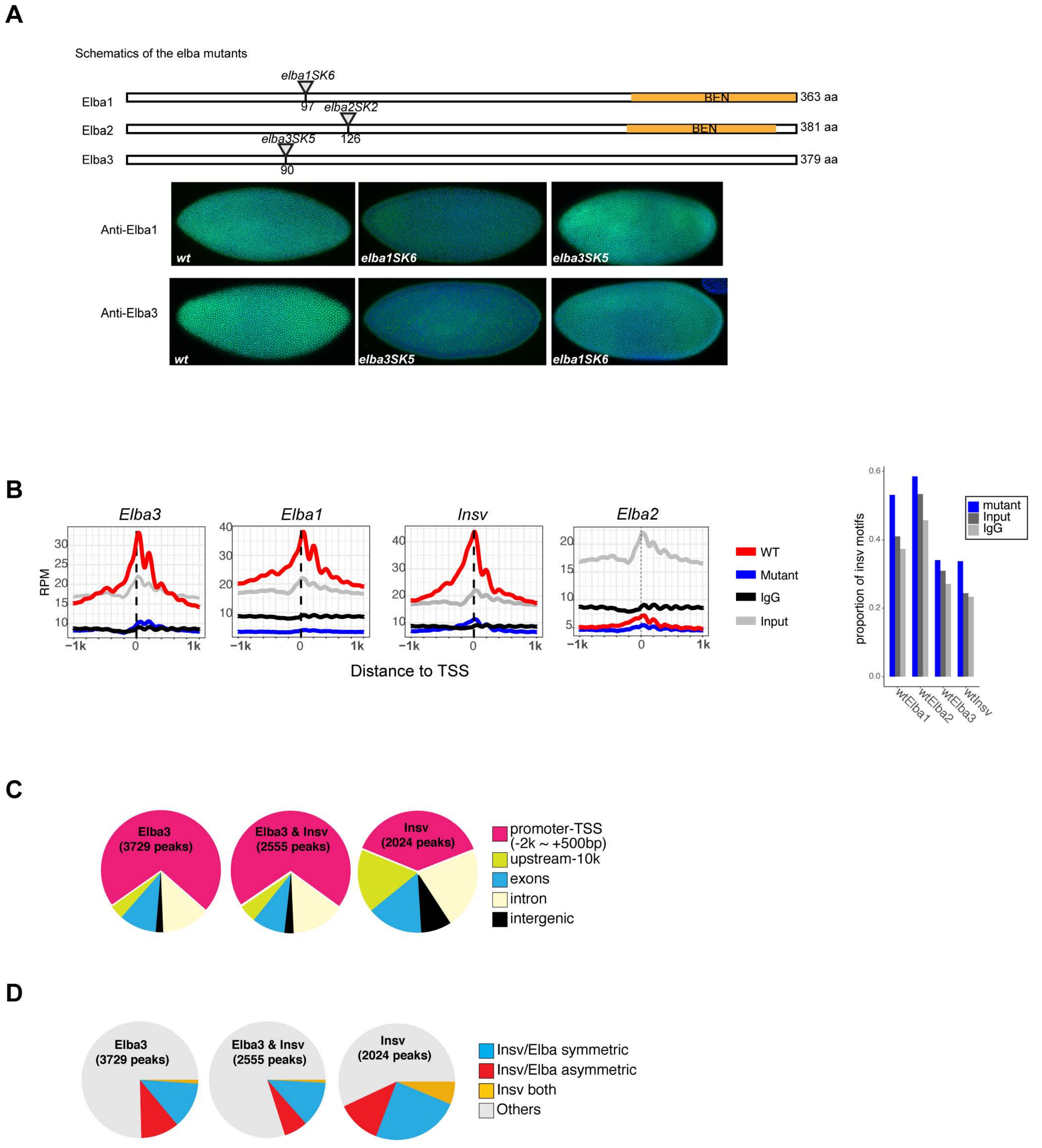
A
Schematics of the elba mutants
elba1SK6
Elba1
97
BEN
363 aa
elba2SK2
Elba2
126
BEN
381 aa
elba3SK5
Elba3
90
379 aa
Anti-Elba1
wt
elba1SK6
elba3SK5
Anti-Elba3
wt
elba3SK5
elba1SK6
B
Elba3
Elba1
Insv
Elba2
RPM
Distance to TSS
WT
Mutant
IgG
Input
proportion of insv motifs
mutant
Input
IgG
wtElba1
wtElba2
wtElba3
wtInsv
C
Elba3 (3729 peaks)
Elba3 & Insv (2555 peaks)
Insv (2024 peaks)
promoter-TSS (-2k ~ +500bp)
upstream-10k
exons
intron
intergenic
D
Elba3 (3729 peaks)
Elba3 & Insv (2555 peaks)
Insv (2024 peaks)
Insv/Elba symmetric
Insv/Elba asymmetric
Insv both
Others

*Ueberschär_Supplemental_Fig.2*

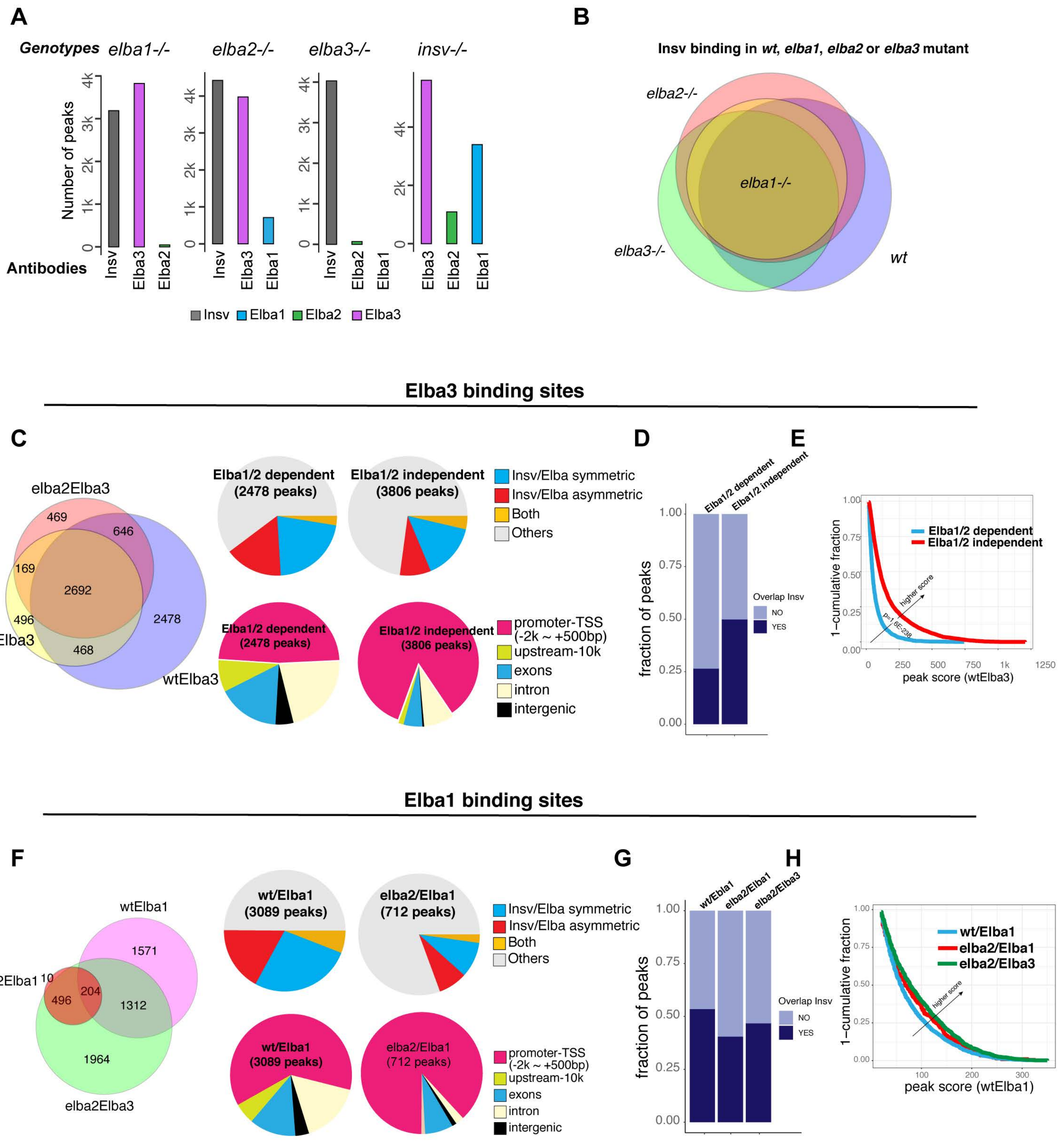

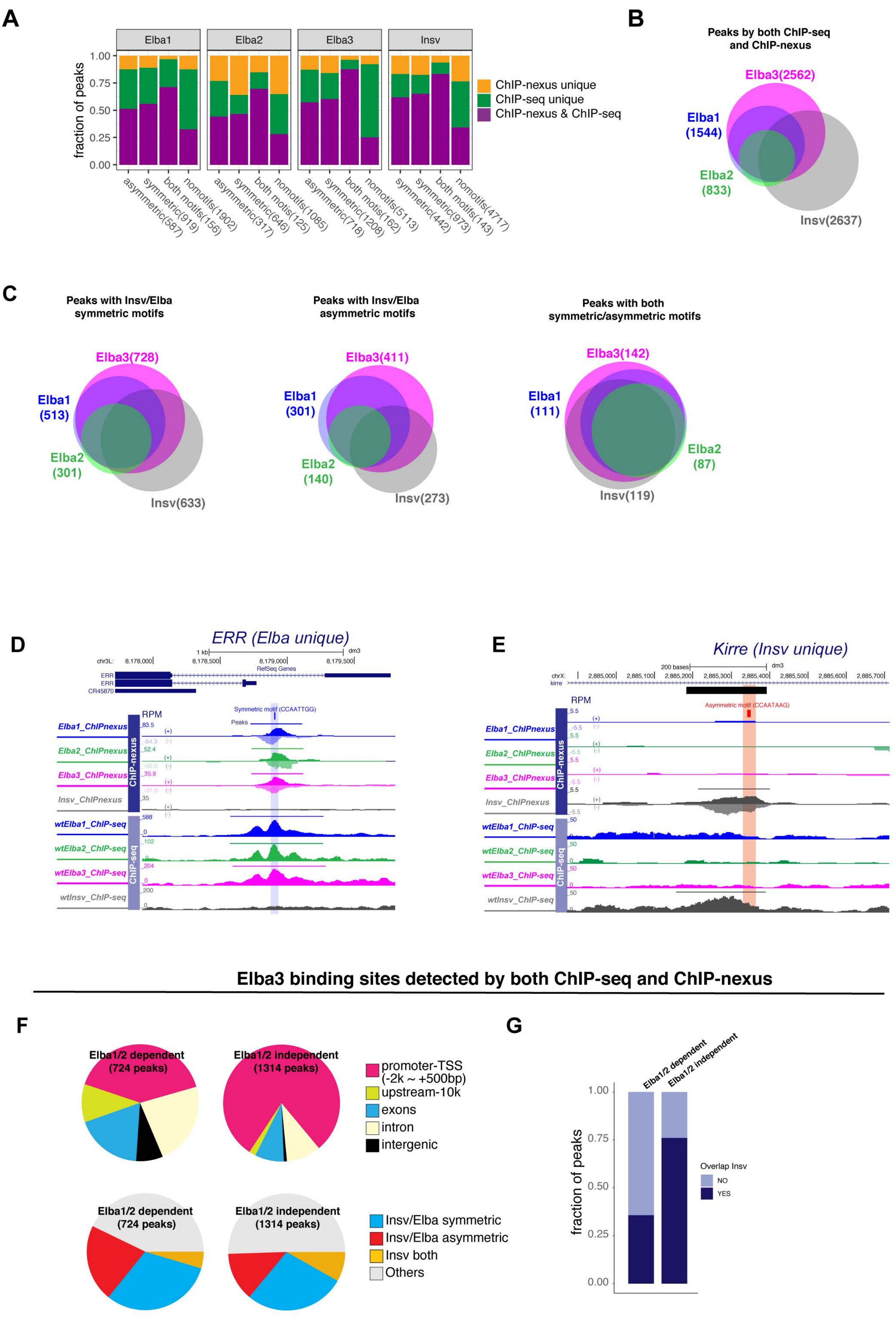
A
Elba1
Elba2
Elba3
Insv
fraction of peaks
1.00
0.75
0.50
0.25
0.00
ChIP-nexus unique
ChIP-seq unique
ChIP-nexus & ChIP-seq
asymmetric(587)
symmetric(919)
both motifs(156)
nomotifs(1902)
asymmetric(317)
symmetric(646)
both motis(125)
nomotifs(1085)
asymmetric(718)
symmetric(1208)
both motis(162)
nomotifs(5113)
symmetric(442)
symmetric(973)
both motifs(143)
nomotifs(4717)
B
Peaks by both ChIP-seq and ChIP-nexus
Elba3(2562)
Elba1 (1544)
Elba2 (833)
Insv(2637)
C
Peaks with Insv/Elba symmetric motifs
Elba3(728)
Elba1 (513)
Elba2 (301)
Insv(633)
Peaks with Insv/Elba asymmetric motifs
Elba3(411)
Elba1 (301)
Elba2 (140)
Insv(273)
Peaks with both symmetric/asymmetric motifs
Elba3(142)
Elba1 (111)
Elba2 (87)
Insv(119)
D
ERR (Elba unique)
Symmetric motif (CCAATTGG)
Peaks
RPM
Elba1_ChIPnexus
Elba2_ChIPnexus
Elba3_ChIPnexus
Insv_ChIPnexus
wtElba1_ChIP-seq
wtElba2_ChIP-seq
wtElba3_ChIP-seq
wtInsv_ChIP-seq
ChIP-nexus
ChIP-seq
E
Kirre (Insv unique)
Asymmetric motif (CCAATAAG)
RPM
Elba1_ChIPnexus
Elba2_ChIPnexus
Elba3_ChIPnexus
Insv_ChIPnexus
wtElba1_ChIP-seq
wtElba2_ChIP-seq
wtElba3_ChIP-seq
wtInsv_ChIP-seq
ChIP-nexus
ChIP-seq
Elba3 binding sites detected by both ChIP-seq and ChIP-nexus
F
Elba1/2 dependent (724 peaks)
Elba1/2 independent (1314 peaks)
promoter-TSS (-2k ~ +500bp)
upstream-10k
exons
intron
intergenic
Elba1/2 dependent (724 peaks)
Elba1/2 independent (1314 peaks)
Insv/Elba symmetric
Insv/Elba asymmetric
Insv both
Others
G
Elba1/2 dependent
Elba1/2 independent
fraction of peaks
1.00
0.75
0.50
0.25
0.00
Overlap Insv
NO
YES

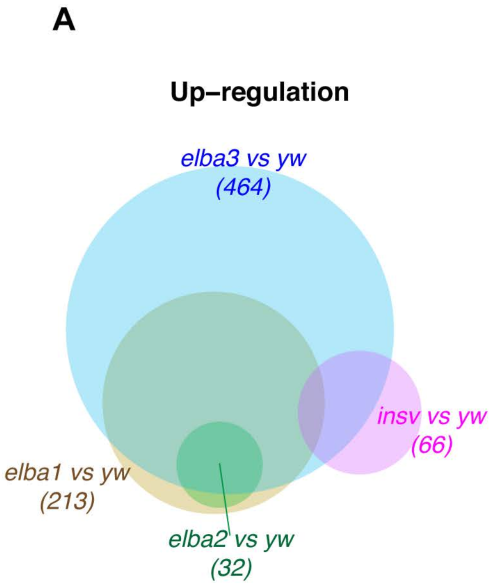
A
Up–regulation
elba3 vs yw
(464)
insv vs yw
(66)
elba1 vs yw
(213)
elba2 vs yw
(32)

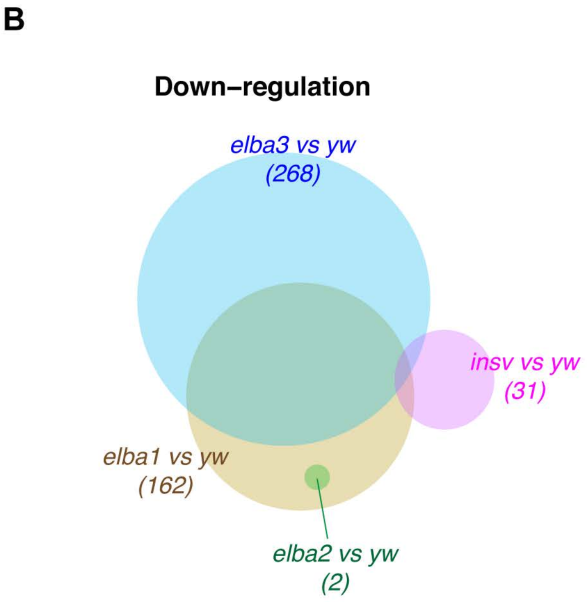
B
Down–regulation
elba3 vs yw
(268)
insv vs yw
(31)
elba1 vs yw
(162)
elba2 vs yw
(2)

A

| Genetic interaction of elba and insulator mutants - Survival | |
|---|---|
| **GAGA factor/Trl** | |
| elba1/elba1;FRT Trl R85/TM3Sb | viable |
| elba2/elba2;FRT Trl R85/TM3Sb | lethal |
| elba2/elba2;FRT Trl R85/pBac(insvelba2) | viable |
| elba2/elba2;Trl 13C/TM3Sb | viable |
| elba3/elba3;FRT Trl R85/TM3Sb | lethal |
| elba3/elba3;Trl 13C/TM3Sb | viable |
| insv/insv;FRT Trl R85/TM3Sb | viable |
| **CP190** | |
| elba1/elba1;CP190P11/TM3Sb | lethal |
| elba2/elba2;CP190P11/TM3Sb | viable |
| elba3/elba3;CP190P11/TM3Sb | lethal |
| insv/insv;CP190P11/TM3Sb | viable |

*Ueberschär_Supplemental_Fig.6*

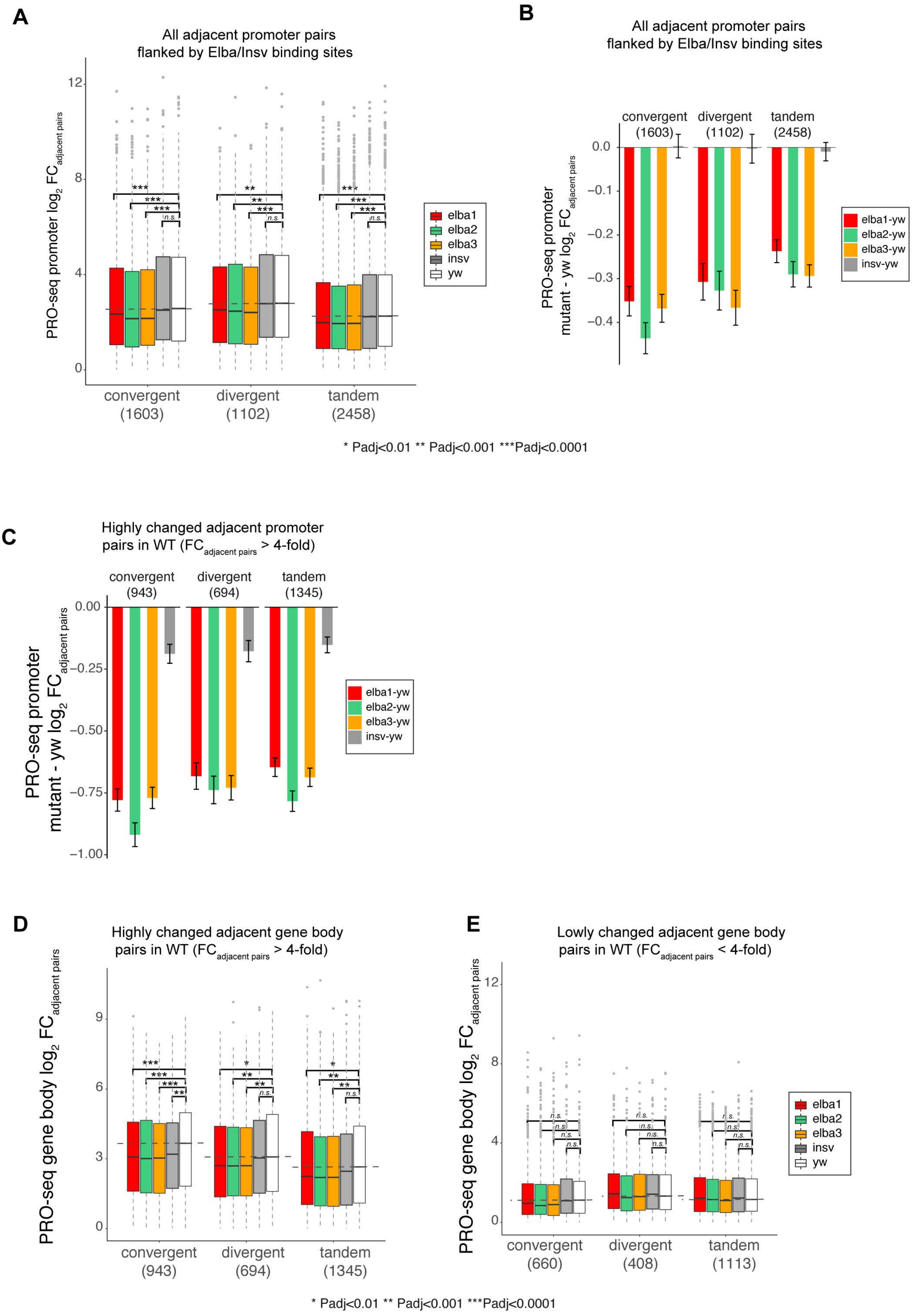

A

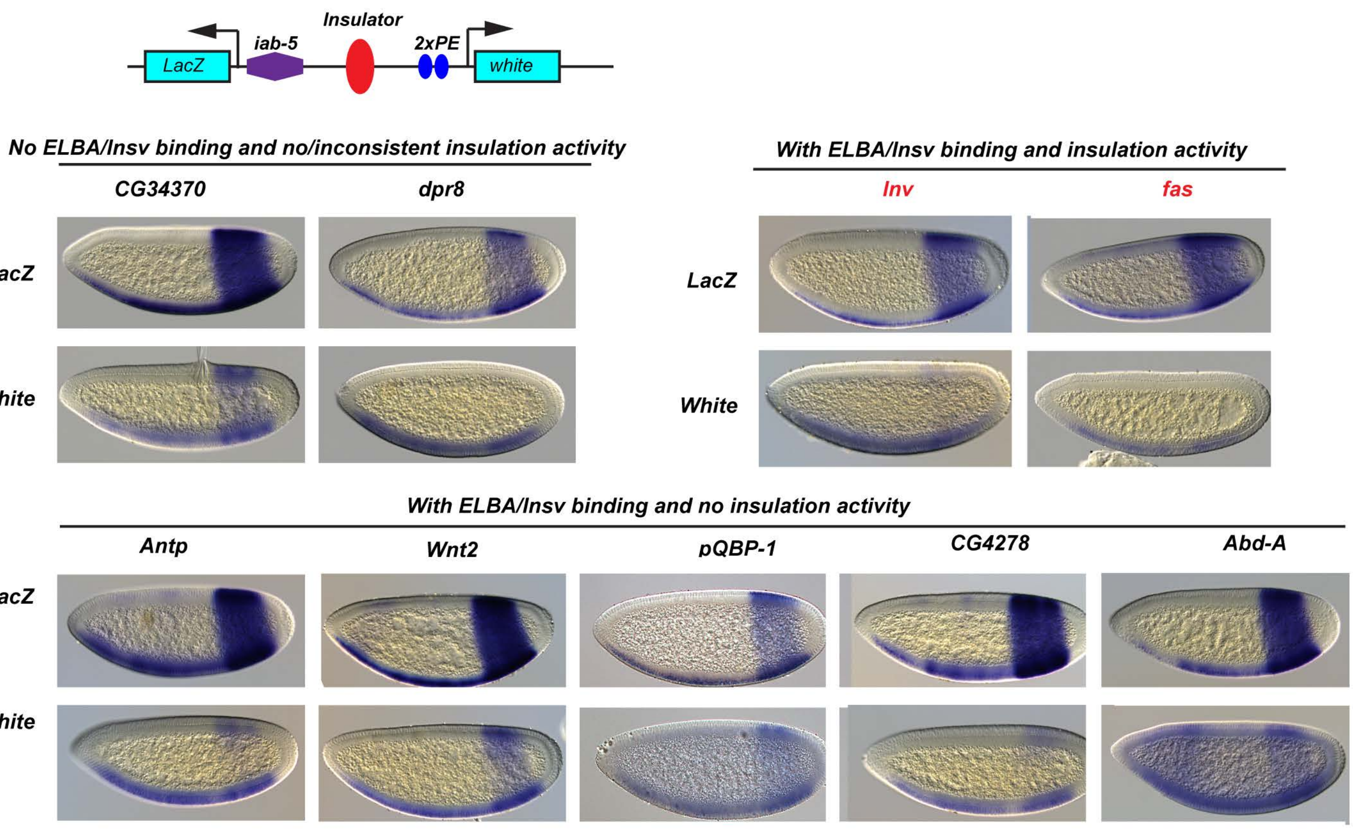

B

| | Insulation | multiple l es | Elba-bound | Insv-bound | Elba motif |
|---|---|---|---|---|---|
| µMAR | N | n | | | |
| Abd-A | N | n | + | + | |
| CG4278 | N | n | + | + | CCAATAAG |
| Antp | N | n | Elba3 | + | - |
| PQBP-1 | N | n | Weak Elba3 | + | - |
| Wnt2 | N | n | + | + | - |
| inv | Y | y | + | + | - |
| fas | y | y | weak | + | - |
| Lasp | Y directional | y | Elba3, weak Elba1 | + | - |
| wg | Y | y | ++ | + | CCAATAAG |
| CG42368 | y | y | + | + | - |
| Parp | Y | y | + | ++ | CTTATTGGT CTTATTGG |
| CG34370 | N | n | - | - | - |
| CG32333 | N | n | - | - | - |
| dpr8 | y | Inconsistent | - | - | - |
